# A histone methyltransferase-independent function of PRC2 controls small RNA dynamics during programmed DNA elimination in *Paramecium*

**DOI:** 10.1101/2023.07.04.547679

**Authors:** Caridad Miró-Pina, Olivia Charmant, Marina Giovannetti, Augustin de Vanssay, Andrea Frapporti, Adeline Humbert, Maoussi Lhuillier-Akakpo, Guillaume Chevreux, Olivier Arnaiz, Sandra Duharcourt

## Abstract

To limit transposable element (TE) mobilization, most eukaryotes have evolved small RNAs to silence TE activity via homology-dependent mechanisms. Small RNAs, 20-30 nucleotides in length, bind to PIWI proteins and guide them to nascent transcripts by sequence complementarity, triggering the recruitment of histone methyltransferase enzymes on chromatin to repress the transcriptional activity of TEs and other repeats. In the ciliate *Paramecium tetraurelia*, 25-nt scnRNAs corresponding to TEs recruit Polycomb Repressive Complex 2 (PRC2), and trigger their elimination during the formation of the somatic nucleus. Here, we sequenced sRNAs during the entire sexual cycle with unprecedented resolution. Our data confirmed that scnRNAs are produced from the entire germline genome, from TEs and non-TE sequences, during meiosis. Non-TE scnRNAs are selectively degraded, which results in the specific selection of TE-scnRNAs. We demonstrate that PRC2 is essential for the selective degradation of non-TE-scnRNAs, independently of its histone methyltransferase activity. We further show that the PRC2 cofactor Rf4 is required for the physical interaction between the scnRNA-binding protein Ptiwi09 and the zinc finger protein Gtsf1, pointing to an architectural role of PRC2 in scnRNA degradation.

## Introduction

Small silencing RNAs bound to Argonaute/Piwi proteins target RNAs for degradation. Conversely, endogenous RNAs are able to trigger degradation of their cognate small RNA. Instances of target-directed degradation of microRNAs have been reported in mammalian cells, *Caenorhabditis elegans* and *Drosophila* (1). In ciliates, a different class of small RNAs, the scnRNAs, that resemble the metazoan Piwi-interacting RNAs (piRNAs), undergo selective degradation during developmentally programmed DNA elimination (2–6).

In the ciliate *Paramecium tetraurelia*, two types of nuclei coexist in the same cell: the germline micronucleus (MIC) and the somatic macronucleus (MAC). During the self-fertilization process of autogamy, the MIC undergoes meiosis and karyogamy to produce the zygotic nucleus. New MICs and new MACs develop from mitotic products of the zygotic nucleus (Figure 1) (7). The MAC contains a reduced genome (72 Mb) compared to the MIC genome (108 Mb) as a result of massive and reproducible genome elimination (∼30 Mb) that occurs at each sexual cycle (8–10). Eliminated sequences include 45, 000 Internal Eliminated Sequences (IESs) that are remnants of transposable elements scattered throughout the genome (10). Other Eliminated Sequences (OES) (11) correspond to large regions comprising repeats such as transposable elements or satellites (9).

**Figure 1.**
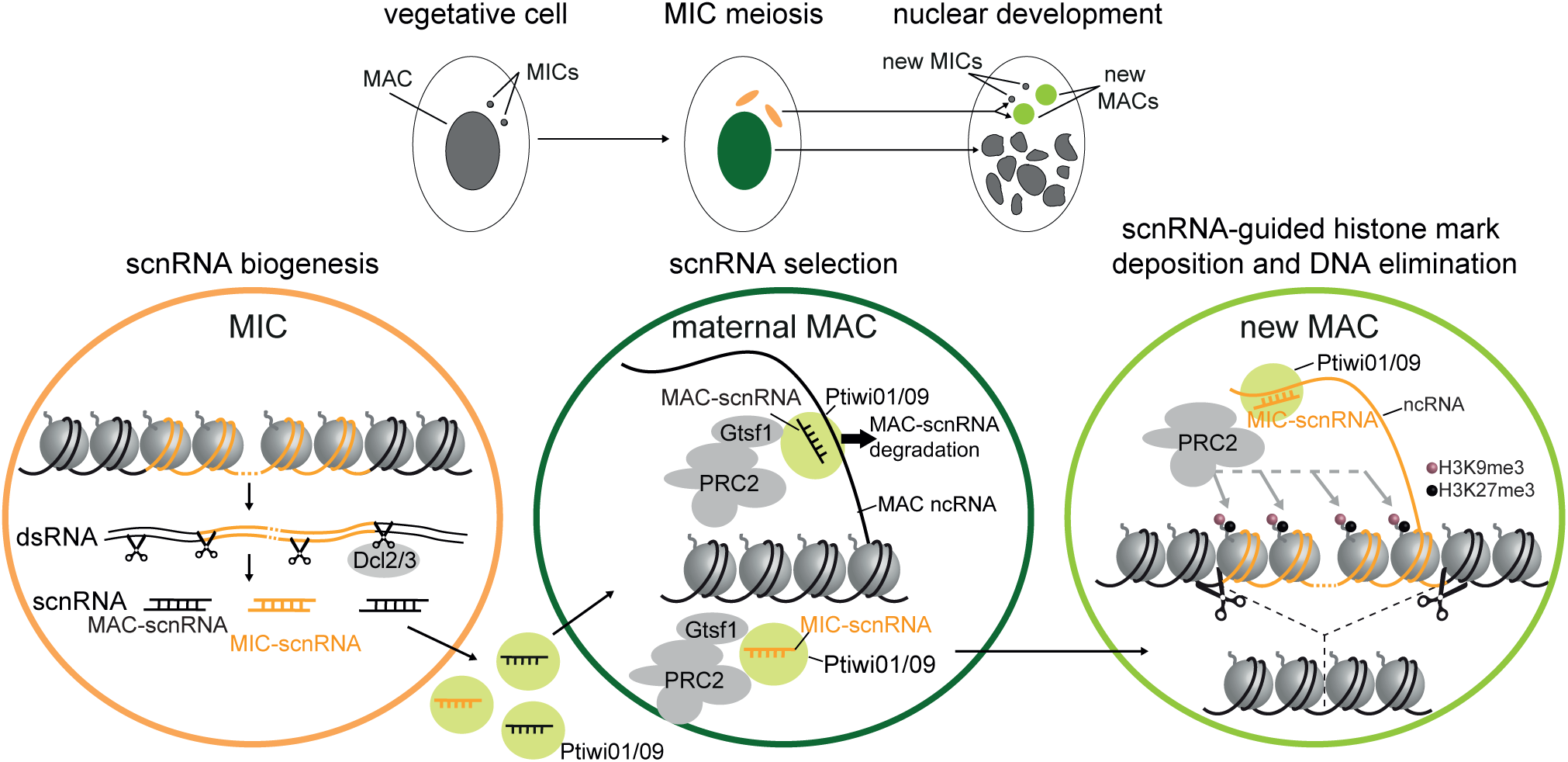
The genome scanning model. The progression of nuclear developmental events during autogamy is represented above, and below, the molecular steps of the scanning model. (i) scnRNA biogenesis. During meiosis, 25-nt scnRNA are produced by the Dicer-like proteins Dcl2/3 from the whole MIC genome (MAC-destined sequences in black, MIC-specific sequences in orange) and are loaded onto Ptiwi01/09. (ii) scnRNA selection. Ptiwi–scnRNA complexes are transported to the maternal MAC. Homologous pairing between scnRNAs and MAC non-coding RNAs (ncRNA) leads to the degradation of MAC-scnRNAs (black) mediated by the zinc finger protein Gtsf1, and selection of MIC-specific scnRNAs (orange) (iii) scnRNA-guided histone mark deposition and DNA elimination. scnRNAs are transported to the new developing MAC, where they recognize homologous sequences by base-pairing interactions with nascent ncRNAs. Interaction between Ptiwi09 and PRC2 guides the deposition of H3K9me3 and H3K27me3 on MIC-specific sequences. These histone marks would guide the recruitment of the excision complex, allowing elimination of MIC-specific sequences (orange).

The sites of DNA elimination do not seem to be specified by the DNA sequence in the MIC genome because, beyond an invariable TA at IES extremities, no strictly conserved sequence motif has been identified in or around IESs. Instead, DNA elimination is epigenetically controlled by the maternal MAC. For a subset of IESs, the artificial introduction of an IES in the maternal MAC was shown to inhibit the elimination of the zygotic IES in the new MAC of the sexual progeny (12, 13). To explain the sequence specificity of this maternal control, a model has been proposed whereby a crosstalk between the maternal MAC and the new MAC is achieved by a genome-wide sequence comparison mediated by RNA molecules (3). 25-nt long scnRNAs produced from germline MIC transcripts by the Dicer-like proteins Dcl2 and Dcl3 (2, 4, 6) are bound to Ptiwi01/09 proteins and transported to the maternal MAC (14). It is believed that scnRNAs are then sorted out by pairing interactions with nascent non-coding RNAs produced by the MAC genome (Figure 1) (3, 15, 16). Non-coding RNAs are thought to trigger the degradation of their cognate small RNA (MAC-scnRNAs), while scnRNAs corresponding to MIC-specific sequences (hereafter called MIC-scnRNAs), which by definition cannot pair with MAC transcripts, are thought to be retained. This selective degradation of MAC-scnRNAs would thus result in the specific selection of the subpopulation corresponding to MIC-scnRNAs. In support of this model, the requirement of complementary MAC transcripts for scnRNA selection has been directly demonstrated in *Paramecium* (16).

The complexes formed by MIC-scnRNAs bound to Ptiwi01/09 are transported from the maternal MAC to the new MAC. In a mechanism very similar to that described in fungi and metazoans (17), scnRNA/Piwi complexes pair to complementary nascent transcripts (18) and guide heterochromatin formation in the new MAC, thereby providing specificity despite the lack of conserved motifs on eliminated sequences (Figure 1). A physical interaction between Ptiwi09 and the Polycomb Repressive Complex 2 (PRC2-Ezl1) was recently shown to mediate the establishment of heterochromatin at scnRNA-targeted regions in the new MAC (Figure 1) (19, 20). PRC2-Ezl1 comprises a core complex and additional cofactors; the catalytic subunit Ezl1 trimethylates histone H3 on lysines 27 and 9 (H3K27me3 and H3K9me3) in the new MAC (21). Deposition of these repressive marks is essential for the elimination of transposable elements and of 70% of IESs (21, 22). Both H3K27me3 and H3K9me3 marks are associated with transposable elements (21). A second class of sRNAs called iesRNAs (26-29 nt), produced from excised IES transcripts by a distinct Dicer protein, Dcl5, and bound to Ptiwi10/11, is reported to enhance the excision of a subset of IESs (4, 23). Both classes of sRNAs appear to act synergistically in the new MAC. Indeed, RNAi-mediated co-depletion of the 3 Dicer proteins Dcl2, Dcl3 and Dcl5 inhibited the excision of 50% of IESs, while depletion of Dcl5 alone or co-depletion of Dcl2 and Dcl3 inhibited the excision of a much smaller subset of IESs (5 and 7% of IESs, respectively) (4, 22).

Although scnRNA selection is a key process in the regulation of DNA elimination, its molecular mechanism remains elusive. We and others recently identified the *Paramecium* zinc-finger protein Gtsf1, which associates with Ptiwi09, as essential for the degradation of MAC-scnRNAs (24, 25). To further understand the mechanism of scnRNA selection, we sequenced small RNAs during the entire sexual cycle of autogamy with unprecedented resolution. Our results provide a comprehensive picture of the dynamics of scnRNAs and of their timing of production. Thanks to these data we confirm recent reports that PRC2 and its cofactors are required for scnRNA selection, thus making an essential contribution to DNA elimination earlier than previously realized. We further show that scnRNA selection relies on a histone methyltransferase independent function of PRC2, demonstrating a novel mode of action for PRC2 in small RNA biology. Our data point to an archirectural role of PRC2 in bringing together Ptiwi09 and Gtsf1.

## MATERIALS AND METHODS

### *Paramecium* strains, cultivation, and autogamy

Experiments were carried out with the entirely homozygous strain 51 of *P. tetraurelia* (wild-type or mutants nd7-1). We used two Rdr mutant strains isolated in (26). The rdr1-5.28 line corresponds to a putative null allele for Rdr1, containing a frameshift mutation leading to a premature stop codon upstream of the catalytic domain. The rdr2–1.24 line is a strong hypomorphic allele that retains an intragenic IES (located downstream of the conserved RdRP domain region), due to a T-to-A mutation in one of the boundaries required for excision during macronuclear development. Cells were grown in wheat grass powder (WGP) infusion medium bacterized the day before use with *Klebsiella pneumoniae*, unless otherwise stated, and supplemented with 0.8 μg/mL β-sitosterol. Cultivation and autogamy (self-fertilization process) were carried out at 27 °C as described (27, 28).

### Gene silencing experiments

For T7Pol-driven dsRNA production in silencing experiments plasmids L4440 with fragments of each gene and *Escherichia coli* strain HT115 DE3 were used. Sequences used for silencing of *ICL7a, EZL1, SUZ12.like*, *RF2*, *RF4*, *EAP1*, *DCL2*, *DCL3, GTSF1* and *EMA1* were segments 1-580 of PTET.51.1.G0700039 (*ICL7a*); 989-1501 of PTET.51.1.G1740049 (*EZL1*); 590-946 of PTET.51.1.G0190277 (*SUZ12.like*); 1514-1916 of PTET.51.1.G1190062 (*RF2*); 623-1038 of PTET.51.1.G0570234 (*RF4*); 298-486 of PTET.51.1.G1310069 (*EAP1*); 434-1032 of PTET.51.1.G0210241 (*DCL2*); 630-1468 of PTET.51.1.G0990073 (*DCL3*); and 249–479 of PTET.51.1.G0490019 (*GTSF1*) (pOC19) as previously published (2, 19, 22, 24, 29). For silencing of *PTIWI01/09*, segments 51-439 of PTET.51.1.G0660118 (*PTIWI09*) and 41-441 of PTET.51.1.G0710112 (*PTIWI01*) (pMG11) were cloned between T7 promoters in plasmid L4440. Preparation of silencing medium and RNAi during autogamy were performed as described in (30).

### Transformation with tagged fusion transgenes

The *GFP-PTIWI09*, *GFP-EZL1_H526A_*, *3XFLAG-HA-EZL1*, *3XFLAG-HA-PTIWI09* (pAH30)*, 3XFLAG-HA-GTSF1* (pOC17) transgenes were previously described (21, 24, 31). For the construction of the *3XFLAG-HA-PTIWI09_DA_* mutant transgene (pMG18) the aspartic acid-coding triplet GCT from nucleotide 1674 to 1676 (D541) of PTET.51.1.G0660118 was replaced by GAT (alanine). pAH30 and pMG18 are RNAi-resistant. For the construction of *H3_K27M_* mutant transgenes, the lysine-coding triplet AAA from nucleotide 85 to 87 (K27) was replaced by ATG (methionine) in the *H3P1* (PTET.51.1.G1450006) and in the *H3P3* (PTET.51.1.G0270094) genes. Each gene was expressed under the control of their transcription signals (promoter and 3’UTR). The *H3P1* (PTET.51.1.G1450006) (pAH2) transcription signals contains 150-bp upstream and 192-bp downstream of its open reading frame. The *H3P3* (PTET.51.1.G0270094) (pAH11) transcription signals contains 89-bp upstream and 147-bp downstream of its open reading frame. For the *H3P3_K27M_-GFP* construct (pAH13), a GFP tag codon-optimized for the *P. tetraurelia* genetic code (same as *GFP-PTIWI09* and *GFP-EZL1_H526A_*) was added to the 3’ end of the gene. Plasmids carrying the transgenes were linearized with restriction enzymes and microinjected into the MAC of vegetative 51 cells in the macronucleus.

To measure the copy number of the transgene per haploid genome (cphg), the transgene copy number is normalized to the endogenous *EZL1* gene copy number by qPCR. qPCR was performed using LightCycler® 480 SYBR Green I Master (catalog number 04 707 516 001, Roche) on the Light Cycler 480 system (Roche). qPCR amplification was done with primers listed in Table S1.

### Production of sexual progeny

Lethality after autogamy was assessed by transferring 30–60 individual post-autogamous cells to standard growth medium and measuring their ability to resume vegetative growth. Cells with a functional new MAC were identified as normally growing survivors unable to undergo a novel round of autogamy if starved after ∼8 divisions.

### Indirect immunofluorescence and GFP localization

For immunofluorescence, cells were fixed as previously described (21) and incubated overnight at room temperature with primary antibodies as follows: rabbit anti-H3K9me3 (1:200) (21), rabbit anti-H3K27me3 (1:1000) (21) and mouse anti-FLAG (1:200) (Thermo Fisher Scientific Cat#MA1-91878; RRID: AB_1957945). Cells were labeled with Alexa Fluor 568-conjugated goat anti-rabbit IgG (Thermo Fisher Scientific Cat#A-11036; RRID: AB_10563566) or Alexa Fluor 568-conjugated goat anti-mouse IgG (Thermo Fisher Scientific Cat#A-11031; RRID: AB_144696) at 1:500 for 1 h, stained with 1 μg/mL Hoechst for 5–10 min and finally mounted in Citifluor AF2 glycerol solution (Biovalley Cat#AF2-25). For GFP detection, cells were fixed for 15 minutes in PHEM 1X, 4 % formaldehyde, 4 % sucrose at room temperature and washed in Tris buffered saline-Tween 20 0.1 % (TBST). Hoechst staining was performed with 1 μg/mL Hoechst for 5–10 min. Cells were washed in Tris buffered saline-Tween 20 0.1 % (TBST) and mounted in Citifluor AF2 glycerol solution. Images were acquired using a Zeiss LSM 980 laser-scanning confocal microscope and a Plan-Apochromat 63 × /1.40 oil DIC M27 objective. Z-series were performed with Z-steps of 0.35 μm.

### Immunoprecipitation

*Paramecium* nuclear protein extracts were performed as previously described (21). 10_6_ autogamous cells (T0) were lysed with a Potter-Elvehjem homogenizer in 3 volumes of lysis buffer (10 mM Tris pH 6.8, 10 mM MgCl2, 0.2 % Nonidet P-40, 1 mM PMSF, 4 mM benzamidine, 1x Complete EDTA-free Protease Inhibitor Cocktail tablets (Roche)). The nuclei-containing pellet was collected by centrifugation and washed with the addition of 2.5 volumes of washing solution (0.25 M sucrose, 10 mM MgCl2, 10 mM Tris pH 7.4, 1 mM PMSF, 4 mM benzamidine, 1x Complete EDTA-free Protease Inhibitor Cocktail tablets (Roche)). The pellet was incubated in 1 volume of nuclear extraction buffer 2 × (100 mM Hepes pH 7.8, 100 mM KCl, 300 mM NaCl, 0.2 mM EDTA, 20 % Glycerol, 2 mM DTT, 0.02 % Nonidet P-40, 2 mM PMSF, 2x Complete EDTA-free Protease Inhibitor Cocktail tablets (Roche)) for 1 hour at 4°C. The salt-extractable nuclear fraction at 150 mM NaCl was recovered following centrifugation for 3 minutes at 10, 000 g at 4°C. Nuclear extracts were incubated overnight at 4°C with 150 µl anti-FLAG M2 magnetic beads (M8823, Sigma) that were pre-washed with 1 mL TEGN buffer (20 mM Tris pH 8, 0.1 mM EDTA, 10 % Glycerol, 150 mM NaCl, 0.01 % Nonidet P-40). Beads were washed five times with TEGN buffer and eluted with 3xFLAG peptide (F4799, Sigma-Aldrich) (45 µL) and the same volume of TEGN buffer at 4°C for 5 hours. RNase I treatment in anti-FLAG Gtsf1 IP was performed as described in (19) except for the use of 2 mM Ribonucleoside Vanadyl Complex (NEB) instead of 4 mM benzamidine. After incubation of half of the beads with 200 U of RNase I (Thermo Fisher Scientific) for 5h at 4°C, beads were washed three times with TEGN buffer and boiled in Laemmli sample buffer.

### Mass spectrometry

#### Sample preparation

Gel plugs were washed with a destaining solution of ACN/NH_4_HCO_3_ 50 mM (50/50), then reduced with Tris(2-Carboxyethyl) Phosphine Hydrochloride (TCEP-HCl) 10 mM and alkylated with S-Methyl methanethiosulphonate (MMTS) 20 mM. Proteins were then in gel-digested overnight at 37 °C with 20 μL of 50 mM NH_4_HCO_3_ buffer containing 1 µg of sequencing-grade trypsin/Lys C mix (Promega). The digested peptides were loaded and desalted on evotips provided by Evosep (Odense, Denmark) according to manufacturer’s procedure before LC-MS/MS analysis.

#### LC-MS/MS acquisition

Samples were analyzed on a timsTOF Pro 2 mass spectrometer (Bruker Daltonics, Bremen, Germany) coupled to an Evosep one system (Evosep, Odense, Denmark) operating with the 30SPD method developed by the manufacturer. Briefly, the method is based on a 44-min gradient and a total cycle time of 48 min with a C18 analytical column (0.15 x 150 mm, 1.9µm beads, ref EV-1106) equilibrated at 40°C and operated at a flow rate of 500 nL/min. H_2_O/0.1 % FA was used as solvent A and ACN/ 0.1 % FA as solvent B. The timsTOF Pro 2 was operated in Data Dependent Analysis-Parallel Accumulation Serial Fragmentation (DDA-PASEF) mode over a 1.3 sec cycle time. Mass spectra for MS and MS/MS scans were recorded between 100 and 1700 *m/z*.

#### Data analysis

MS raw files were processed using PEAKS Online 11 (build 1.9, Bioinformatics Solutions Inc.). Data were searched against the *Paramecium tetraurelia* database downloaded from ParameciumDB website (parameciumtetraurelia_mac_51_annotation_v2.0.protein.fasta, 40460 entries). Parent mass tolerance was set to 20 ppm, with fragment mass tolerance to 0.05 Da. Specific tryptic cleavages were selected and a maximum of 2 missed cleavages were allowed. The following post-translational modifications were considered for identification: Oxidation (M), Deamidation (NQ), Acetylation (Protein N-term) as variable and Beta-methylthiolation (C) as fixed. Identifications were filtered based on a 1% FDR (False Discovery Rate) threshold at both peptide and protein group levels. Label free quantification was performed using the PEAKS Online 11 quantification module, allowing a mass tolerance of 5 ppm, a CCS error tolerance of 0.01 and a 0.25-min retention time shift tolerance for match between runs. Protein abundance was inferred using the top N peptide method and TIC was used for normalization. Multivariate statistics on protein measurements were performed using Qlucore Omics Explorer 3.9 (Qlucore AB, Lund, *SWEDEN*). A positive threshold value of 1 was set to allow a log2 transformation of abundance data for normalization *i.e.* all abundance data values below the threshold are replaced by 1 before transformation. The transformed data were finally used for statistical analysis *i.e.* the evaluation of differentially present proteins between two groups using a bilateral Student’s t-test.

### Western blot and silver staining

Electrophoresis and blotting were carried out according to standard procedures. Samples were run on 10% Tris-Glycine polyacrylamide gels (Bio-Rad). Blotting was performed for 1 hour at 100 V using a nitrocellulose membrane (GE10600002, Merck). FLAG (1:1000) (MAI-91878, Thermo Fisher Scientific), α-tubulin (1:5000) (sc-8035, Santa Cruz), *Paramecium* H3 (1:5000) (21), HA (1:2000) (H6908, Merck) and Ezl1 (1:1000) (19) were used for primary antibodies. Secondary horseradish peroxidase-conjugated anti-mouse or anti-rabbit IgG antibodies (Promega) were used at 1:2500 dilution followed by detection by ECL. Alternatively, secondary fluorescent anti-mouse (DyLight 800, BioRad) or anti-rabbit (StarBright700) IgG antibodies were used at 1:2000 dilution.

For silver staining, samples were run on 4-20% Tris-Glycine extended polyacrylamide gels (Cat #4568095, BioRad) for 20 minutes at 200 V and the fast protocol of the SilverQuest Silver Staining Kit (Thermo) was used according the manufacturer’s instructions.

### Small RNA isolation and immunoprecipitation

200 mL of cells were collected and frozen at T0 after the onset of autogamy. FLAG-Ptiwi09 immunoprecipitation was performed as described in (14). Cells were lysed using a dounce homogenizer in 2 mL lysis buffer (50 mM Tris HCL pH 8, 150 mM NaCl, 5 mM MgCl2, 1 mM DTT, 0.5 % sodium deoxycholate, 2 mM vanadyl ribonuclease complex (VRC), 1% Triton X-100, 10 % glycerol, 1x Complete EDTA-free Protease Inhibitor Cocktail tablets (Roche)). 1 mL lysate was used to perform immunoprecipitation with 70 μL of anti-FLAG M2 magnetic beads (M8823, Sigma) (pre-washed 5 times in lysis buffer) overnight at 4°C. The unbound fraction was kept and the beads were washed 5 times with IP buffer (50 mM Tris HCL pH 8, 150 mM NaCl, 5 mM MgCl2, 1 mM DTT, 0.5 % sodium deoxycholate, 2 mM vanadyl ribonuclease complex (VRC), 1 % Triton X-100, 10 % glycerol, 1x Complete EDTA-free Protease Inhibitor Cocktail tablets (Roche)) and resuspended in 1 mL IP buffer. 1/10 of the resuspended beads was kept for western blot analysis. After removal of the supernatant, 300 µl of proteinase K (10 μg/mL) in Tris pH 7.5 (10 mM), EDTA pH 8 (5 mM), 0.5 % SDS is added to the rest of the beads and incubated for 20 minutes at 42 °C. RNA extraction was performed as previously described (21).

### DNA extraction and PCR

DNA samples were prepared from ≤1, 000 autogamous cells using the NucleoSpin Tissue kit (Macherey-Nagel). For PCR analysis of IES excision, PCR amplifications were performed with Expand Long Template Enzyme mix (Expand Long Template PCR system, Roche). Oligos are available in Table S1.

### RNA extraction, RT-PCR and sequencing

RNA samples were extracted from 200–400 mL cultures at 2, 000–4, 000 cells/mL as previously described (21) at different time points during autogamy. For RT-PCR, RNA samples were reverse-transcribed with RevertAid H Minus Reverse Transcriptase (Thermo Scientific) using random hexamer primers (Thermo Scientific) according to the manufacturer’s instructions. It was then followed by PCR amplifications in a final volume of 25 µL, with 10 pmol of each primer, 10 nmol of each dNTP and 2 U of DyNAzyme II DNA polymerase (Thermo Scientific). For sRNA-seq, small RNAs of 15-30 nt (or 10-30 nt for the analysis of the *PTIWI09_DA_* mutant and Ptiwi09 immunoprecipitation) were purified from total RNA on a 15% TBE/Urea gel. Small RNA libraries were constructed using the NEBNext Small RNA kit according to the manufacturer recommendations. The quality of the final libraries was assessed with an Agilent Bioanalyzer, using an Agilent High Sensitivity DNA Kit. Libraries were pooled in equimolar proportions and sequenced using a single read 75 pb run on an Illumina NextSeq500 instrument, using NextSeq 500 High Output 75 cycles kit.

### Bioinformatic analyses

Sequencing data were demultiplexed using CASAVA (v1.8.2) and bcl2fastq2 (v2.18.12). Illumina adapters were removed using cutadapt (v1.12), only keeping reads with a minimal length of ten nucleotides. Paired-end reads were mapped on expected contaminants (mitochondrial genomes, ribosomal DNA, and bacterial genomes). sRNA-seq data were mapped on reference genomes using BWA (v0.7.15 -n 0). sRNA-seq data were analyzed as previously (32). Sequencing metrics are available in Table S2.

### Reference genomes, software and R packages

Sequencing reads were mapped on expected contaminants (mitochondrial genomes, ribosomal DNA, and bacterial genomes) then mapped on *Paramecium* genome references using BWA (v0.7.15) without any mismatches. The *P. tetraurelia* strain 51 reference genomes used in this study were: the MAC (ptetraurelia_mac_51.fa), the MAC+IES (ptetraurelia_mac_51_with_ies.fa) and the MIC (ptetraurelia_mic2.fa) (9, 33). Gene annotation v2.0 (ptetraurelia_mac_51_annotation_v2.0.gff3), IES annotation v1 (internal_eliminated_sequence_PGM_ParTIES.pt_51.gff3) were also used (33, 34). All files are available from the ParameciumDB download section (https://paramecium.i2bc.paris-saclay.fr/download/Paramecium/tetraurelia/51/) (34). R (v4.0.4) packages were used to generate images (ggplot2 v3.3.5; ComplexHeatmap v2.6.2; GenomicRanges v1.42; rtracklayer v1.50).

### sRNA quantification

Small RNA reads were successively mapped on the MAC, MAC+IES and MIC genomes to attribute them to a specific genome compartment: MAC-destined sequence (MDS), Internal Eliminated Sequences (IES) or Other Eliminated Sequences (OES). In the case of a silencing experiment, the 23nt siRNA reads that map to the RNAi targets were removed. Read counts were normalized using the number of sequenced reads using a G+C content < 50%, compatible with a *Paramecium* G+C genomic content (∼27%).

### Scaled genome coverage of scnRNAs

To analyse the evolution of sRNA genome coverage, the sequencing data were sampled to 15M reads between 20 and 30 nt long. After mapping on the MIC genome, only 25-nt sRNA were kept. The coverage of the MIC genome was calculated using bamCoverage (deeptools v2.0) and a bin window of 100 nt. Using this window coverage, the percentage of genome covered by at least 1X was determined for each sample.

### sRNA clusters

Only sRNAs of 23-nt in length mapped on the MAC genome were considered. Sequencing data from all the wild-type autogamy time course were merged to stringently annotate 136 short read clusters (SRC) using the program ShortStack (v3.8.5 --nohp --mincov 1000 --pad 100) (35). The boundaries of these SRCs were manually adjusted and compared to, already published, vegetative SRCs with a minimum overlap of 23-nt (36–38)(see Table S3). The number of 23-nt sRNAs were counted on each SRC, with the information regarding transcription orientation, and normalized by the total number of sequenced sRNAs (20-30 nt with GC < 50%) and their length (RPKM). A SRC and a gene were associated with a minimum overlap of 20 nt. K-means clustering (iter.max=100) was used to define 3 groups of SRCs based on their RPKM coverage during wild-type autogamy time course experiments. siRNA transcription orientation ratio was calculated using counts obtained with SAMTools (samtools view). Unidirectional and bidirectional transcription ratios were calculated only for SRCs overlapping a gene and covered by at least 5 siRNA reads. mRNA RPKM coverage of SRCs was calculated using htseq-count (v0.11.2 stranded=no -mode=intersection-nonempty) on TopHat2 mappings (v2.1.1 --min-intron-length 15 --max-intron-length 100 --read-mismatch 1 --coverage-search) (39).

## Results

### Dynamics of scnRNAs during the sexual cycle

To get a comprehensive picture of scnRNA dynamics, we sequenced small RNA populations at 11 time points covering with unprecedented resolution the entire sexual cycle of autogamy (self-fertilization) (Figure 2). The time course experiments covered the whole duration of pre- and post-zygotic nuclear events, including the time window during which DNA elimination occurs (40). Small RNA populations were mapped with no mismatches against the reference genomes and subdivided in three categories according to their length: 25-nt for scnRNAs; 26-29 nt for iesRNAs and 23-nt for small interfering RNA (siRNAs) (Supplementary Figure S1). (For the analysis of iesRNAs and siRNAs, see Supplementary Text, Figures S1-S2-S3, Tables S3-S4). 25-nt scnRNAs begin to accumulate as soon as meiotic cells are detected in the culture (t-2) and persist until cells undergoing meiosis are no longer detected (T10) (Figure 2A), in line with the idea that scnRNA biogenesis occurs during the meiotic phase.

**Figure 2.**
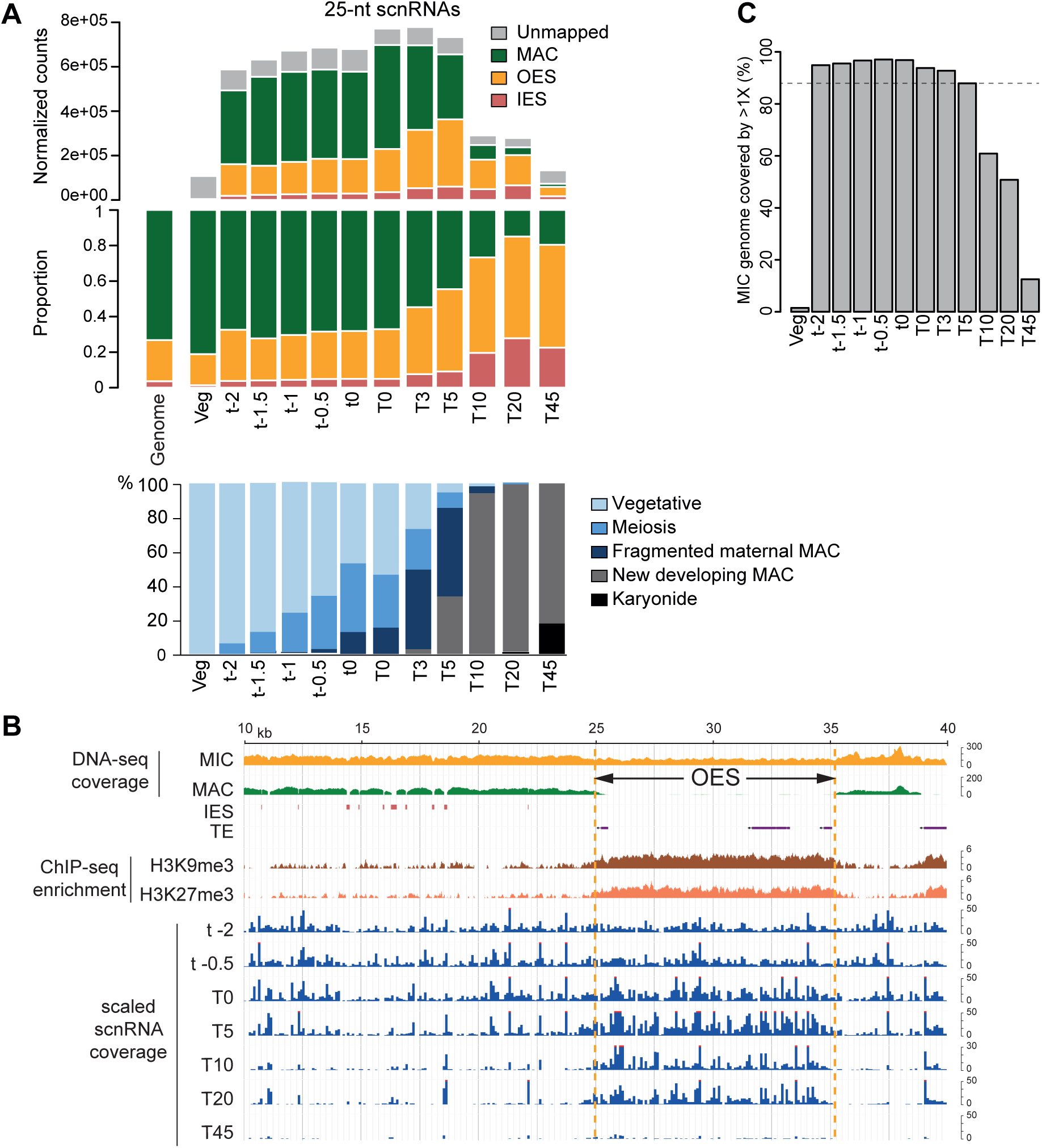
scnRNA dynamics during Paramecium sexual cycle. **A.** Analysis of 25-nt-scnRNA populations in vegetative cells and at different time points during autogamy. Bar plots show the normalized counts, using the total number of sRNAs (see Materials and Method), of 25-nt reads for each sample that that are unmapped or map the MAC genome, IESs, or OES reference sequences (top) and the proportion of 25-nt reads for each catego-ry (middle). Genome: proportion of each category (MAC, OES, IES) in the MIC genome. Progression of autogamy is followed by cytology with Hoechst staining (bottom) and is not synchronous in the cell population (76). Five different morphological states were scored for each autogamy time-course sample: “Vegetative” cells with 2 MICs and a single MAC; “Meiosis” is scored by observation of MIC division at different stages of meiosis I and II; “Fragmented maternal MAC” corresponds to maternal MAC into many small pieces. (Fragments are only completely lost, by dilution, during the first vegetative cell divisions after autogamy); clearly visible “developing new MACs”; at the end of autogamy, the first cellular division distributes the two new MACs to the daughter cells (“karyonides”). Two independent time course (tc1 and tc2) experiments are displayed: tc1 covers veg, t-2 to t0 and tc2 covers T0 to T45. Overlap between tc1 and tc2 is ensured by t0 and T0, which are alike. The numbers correspond to hours before (e.g. t-2) or after T0 (e.g. T5). At least 100 cells were scored for each time point by fluorescence microscopy. B. Representative genomic region depicting 25-nt scnRNAs at different time points of autogamy. DNA-seq coverage (bin=1) for the MIC and the MAC DNA samples on the MIC genome (NODE_1273_length_39901_cov_28.471216 between 10 and 40 kb), -and IES and TE annotations are indicated. ChIP-seq enrichment (deeptools bamCompare bin=5) of H3K9me3 and H3K27me3 compared to the DNA Input sample from Pgm-depleted cells at T50 (ENA accessions ERS6678598, ERS6678600 and ERS6678597) (19) is displayed, indicating that the OES is enriched for both histone marks in the new developing MAC. scnRNA coverage (bin=100) on the MIC genome at different time points of autogamy (T-2 to T45) is indicated at the bottom. To compare time points, the 25-nt sRNA coverages were calculated on equivalent read datasets (15e6 sRNA between 20 and 30-nt long). C. Using the scaled normalized datasets presented in B., the percentage of the MIC genome covered by at least 1X of 25-nt scnRNA was calculated at each time point of autogamy.

Because the cell populations are not synchronous, however (Figure 2A), we cannot determine with certainty which cells produce scnRNAs. We were able to overcome to some degree by taking advantage of an observation we made with a *GFP-PTIWI09* transgene (31). Cells were transformed with this fusion gene, whose expression is under the control of the endogenous sequences up-and downstream of *PTIWI09*, and sexual events were triggered by starvation. GFP fluorescence was detected in the cytoplasm and in the maternal MAC during meiosis I, before it concentrated in the new MAC, as previously reported (31). However, we noted that, in transformants with high copy numbers of the transgene and high GFP-Ptiwi09 protein levels, as analyzed by western blot, 100% of cells were arrested in early meiosis I before each MIC yields four haploid nuclei (Supplementary Figure S1). In these arrested cells, the GFP fluorescence was detected in the meiotic MICs, as well as in the cytoplasm and in the maternal MAC (Supplementary Figure S1). This localization appears different from that of the other transformants that normally progress through autogamy (Supplementary Figure S1), for which GFP was always excluded from the meiotic MIC (31). We used cells expressing GFP-Ptiwi09 arrested in meiosis I as a tool to assess whether scnRNAs can be detected at this stage. Small RNA sequencing experiments showed that 25-nt scnRNAs accumulate in meiosis I-arrested cells expressing GFP-Ptiwi09 (Supplementary Figure S1), suggesting that scnRNAs are produced prior to or during early meiosis I.

As previously reported from the sequencing of small RNAs from three time points (early conjugation time point in (6), and early and late developmental stages during autogamy in (4)), our dataset of 11 time points during autogamy confirmed that scnRNAs are produced from both strands (Supplementary Figure S1) and cover the whole MIC genome (Figure 2B-C). This can be clearly observed when examining the mapping of 25-nt scnRNAs across an individual genomic region (Figure 2B), where MAC sequences, as well MIC sequences (IES or OES), are covered. Using the draft MIC genome assembly (9), we found that >88 % of the MIC genome is covered by at least 1X of normalized sequencing depth, from the moment scnRNAs are detected (t-2) and until T5 (Figure 2C). No bias in scnRNA coverage between IESs and MAC-destined sequences could be observed at the early time points (until T0) (Table S5). We conclude that scnRNAs are produced from the whole MIC genome in *Paramecium tetraurelia*. As autogamy proceeds, we observed an increase in the proportion of 25-nt scnRNAs mapping to OES or IES over scnRNAs mapping to MAC sequences (Figure 2A-B). This increase started from T3 in our time course experiment, and by T20, scnRNAs mostly corresponded to MIC-specific (IES+OES) sequences (Figure 2A). As illustrated for an individual genomic region, scnRNAs cover the entire OES region, that comprises annotated transposable elements, enriched in H3K9me3 and H3K27me3 histone marks in the new MAC, and the IESs scattered in the flanking regions (Figure 2B). Genome-wide analysis of scnRNA coverage showed that it is rather homogeneous whatever the enrichment in histone marks (H3K27me3 and H3K9me3) at T0, as expected. As development proceeds (from T0 to T45), the correlation between scnRNA coverage and histone mark enrichment increases (Supplementary Figure S1), in agreement with the data presented in Figure 2B. As reported for one early and one late developmental time points (4), the enrichment in MIC-scnRNAs is consistent with the predicted effect of scnRNA selection, resulting from the degradation of MAC-scnRNAs, which correspond to regions that are not enriched in histone marks.

### The slicer activity of Ptiwi09 is required for the removal of the passenger strand

Argonaute proteins can carry an endoribonuclease (slicer) activity responsible of the cleavage of the RNA complementary to the guide RNA loaded to the protein. The slicer activity is provided by a conserved catalytic tetrad composed of a DEDX (X is usually H or D) motif in the Piwi domain that adopts an RNaseH fold (41). The scnRNA binding protein Ptiwi09 possesses a DEDH motif, suggesting it may carry a slicer activity (31). Given that MAC-scnRNAs are thought to be degraded upon pairing with complementary RNA molecules (Figure 1), we decided to investigate the role of Ptiwi09 slicer activity in scnRNA selection. We obtained *Paramecium* transformants expressing full-length, 3xFLAG-HA-tagged mutant Ptiwi09 carrying an aspartic acid-to-alanine substitution (D541A) in the first aspartic acid residue of the conserved DEDH motif within the PIWI domain, expected to abolish slicer activity (42–45). Endogenous Ptiwi09 and Ptiwi01 protein expression was specifically depleted through RNA interference while the mutant *3xFLAG-HA-PTIWI09_DA_* transgene was designed to be RNAi-resistant. *In vivo* complementation assay for three independent transformants expressing full-length, 3xFLAG-HA-tagged wild-type Ptiwi09 indicated that the 3xFLAG-HA-Ptiwi09_WT_ (Ptiwi09_WT_) protein is functional and can rescue the lethality and DNA elimination defects associated with Ptiwi01/09 depletion (Figure 3A and Supplementary Figure S4). In contrast, the Ptiwi09_DA_ mutant protein is unable to complement these defects in two independent transformants (Figure 3A and Supplementary Figure S4), indicating that the slicer activity of Ptiwi09 exerts an essential function during sexual events.

**Figure 3.**
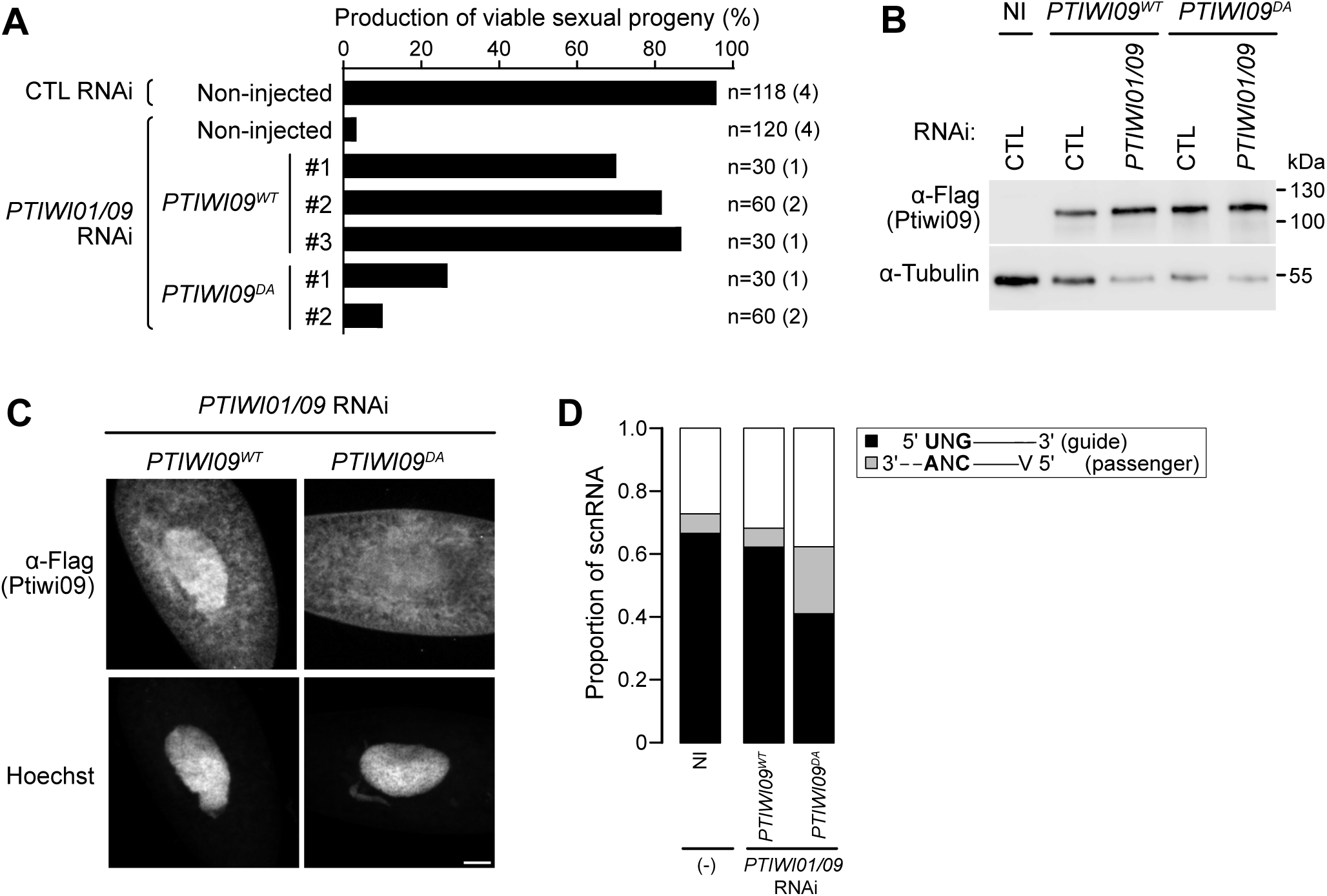
The slicer activity of Ptiwi09 is required for the removal of the passenger strand. A. Production of sexual progeny of non-injected cells and cells transformed with 3XFLAG-HA-PTIWI09 (PTIWI09WT) or 3XFLAG-HA-PTIWI09DA (PTIWI09DA), following PTIWI01/09 or ICL7 (CTL) RNAi. PTIWI09WT transformants #1, #2 and #3 contain respec-tively 0.3 copies of the transgene per haploid genome (cphg), 8.4 cphg and 33 cphg. PTIWI09DA transformants #1 and #2 contain respectively 9.6 cphg and 57 cphg. The total number of cells analyzed for each condition and the number of independent experiments (in parentheses) are indicated. B. Western blot analysis of whole cell extracts from non-injected (NI) cells and cells expressing a 3XFLAG-HA-Ptiwi09 (PTIWI09WT #2) fusion protein or a mutated 3XFLAG-HA-Ptiwi09DA (PTIWI09DA #2) fusion protein, with FLAG antibodies to detect 3XFLAG-HA-Ptiwi09WT or 3XFLAG-HA-Ptiwi09DA and alpha-tubulin antibodies for normalization. Whole cell extracts were done at T0, following PTIWI01/09 or ICL7 (CTL) RNAi. C. FLAG immunostaining at T0 of cells transformed with 3XFLAG-HA-PTIWI09 (PTIWI09WT #2) or 3XFLAG-HA-PTIWI09DA (PTIWI09DA #2), following PTIWI01/09 RNAi. Representative confocal images are displayed. Overlay of Z-projections are presented. Scale bar is 10 µm. D. Analysis of 25-nt scnRNAs at T0 in non-injected (NI) cells and cells transformed with 3XFLAG-HA-PTIWI09 transgenes (tg) (PTIWI09WT #2 or PTIWI09DA #2) in PTIWI01/09 RNAi conditions. Proportion of 25-nt scnRNAs with the 5’-UNG signature, characteristics of the guide strand (black), or those that do not start with a 5’-U (A, G or C: V) and end with a CNANN-3’ signature, characteristics of the passenger strand (grey).

Comparable amounts of Ptiwi09_WT_ and Ptiwi09_DA_ proteins were detected by western blot analysis at T0 (Figure 3B), indicating that the slicer activity is not required for the accumulation of Ptiwi09. Yet the mutant protein displays a different localization compared to the control (Figure 3C; Supplementary Figure S4). Indeed, immunofluorescence experiments with FLAG antibodies indicated that the Ptiwi09_WT_ protein localizes in the cytoplasm and in the maternal MAC at T0, as observed for the GFP-Ptiwi09 fusion protein (31). In contrast, the Ptiwi09_DA_ mutant protein is detected in the cytoplasm throughout autogamy and does not strongly accumulate in the maternal MAC. This indicates that the slicer activity is required for normal Ptiwi09 accumulation in the maternal MAC.

We analyzed the expression of scnRNAs in the absence of the slicer activity of Ptiwi09 and found that scnRNAs accumulate at T0 in the slicer mutant and in wildtype, indicating that the slicer activity is not required for the biogenesis of scnRNAs (Supplementary Figure S4). This is consistent with previous reports showing that the Dcl2 and Dcl3 proteins are responsible for scnRNA production (2, 4, 46). We did not detect shorter small RNAs products that would indicate they are degraded in the slicer mutant (Supplementary Figure S4). scnRNA duplexes are produced by the Dicer-like enzymes with 2-nt 3ʹ overhangs at both ends (46). 5ʹ-UNG scnRNAs, or ‘guide’ strands, are preferentially stabilized (70% of the population), while the complementary CNA of the other strand, or ‘passenger’ strands, are less abundant in the scnRNA population (10%) in wild type conditions (Figure 3D; Supplementary Figure S4). When the slicer Ptiwi09 mutant protein is expressed at T0, we found that the proportion of 5’-UNG scnRNAs is reduced, and that of the passenger strand is increased, whether or not the endogenous Ptiwi01/09 proteins are depleted (Figure 3D; Supplementary Figure S4). We conclude that the slicer activity is used for removal of the passenger strand after loading duplex scnRNAs, and this would allow the formation of functional scnRNA-Ptiwi01/09 complexes. Indeed, catalytically active Argonaute proteins use their slicer activity to cleave the passenger strand of perfect or nearly perfect duplexes generated by Dicer (47–49). Thus, similarly to the *Tetrahymena* Twi1p protein (50), Ptiwi09 slicer activity appears to be essential for strand selection, and precludes the possibility to examine the effects of the slicer acvity in downstream steps.

### PRC2-Ezl1 core complex and its associated cofactors are required for scnRNA selection

ScnRNA selection, which results in the enrichment in MIC-scnRNAs, is thought to occur in the maternal MAC during MIC meiosis (Figure 1). Proteins involved in this process would thus be localized in the maternal MAC at this developmental time. While the PRC2-Ezl1 complex catalyzes deposition of H3K27me3 and H3K9me3 in the new MAC, it is first localized in the maternal MAC during MIC meiosis where it catalyzes solely H3K27me3 (19–22). To assess whether PRC2-Ezl1 is involved in the selection process, we sequenced small RNA populations during autogamy at three different time points (T0; T10 and T35) in cells depleted for the PRC2-Ezl1 core components Ezl1 and Suz12.like and for the accessory proteins Rf2, Eap1 and Rf4 (Figure 4A and Supplementary Figure S5). As previously described upon depletion of PRC2-Ezl1 components (19), the 25-nt scnRNAs are produced from the MIC genome at the beginning of the sexual cycle (T0) with the same proportions of MAC, OES and IES scnRNAs as in control RNAi conditions. In contrast, depletion of the scnRNA biogenesis factors Dcl2/3 leads to the loss of scnRNAs (Figure 4A) (4, 14). However, as development proceeds (T0 and T35), MIC-specific scnRNAs (OES + IES) are not enriched over MAC-specific scnRNAs in PRC2-Ezl1-depleted cells, indicating an absence of MIC-specific scnRNA selection (Figure 4A-B and Supplementary Figure S5). Thus, PRC2-Ezl1 components are broadly required for MAC-scnRNA degradation. This result is in agreement with a previous study showing scnRNA selection defects at a late developmental stage during autogamy upon depletion of the PRC2 core subunit Caf1 (51), as well as a more recent report similarly implicating Ezl1 and Rf4 (20). In cells depleted for Ezl1, Suz12.like, Caf1 and Rf2, the PRC2-Ezl1 complex collapses (19, 20). Therefore, we conclude that PRC2-Ezl1 function in scnRNA selection requires the integrity of the complex.

**Figure 4.**
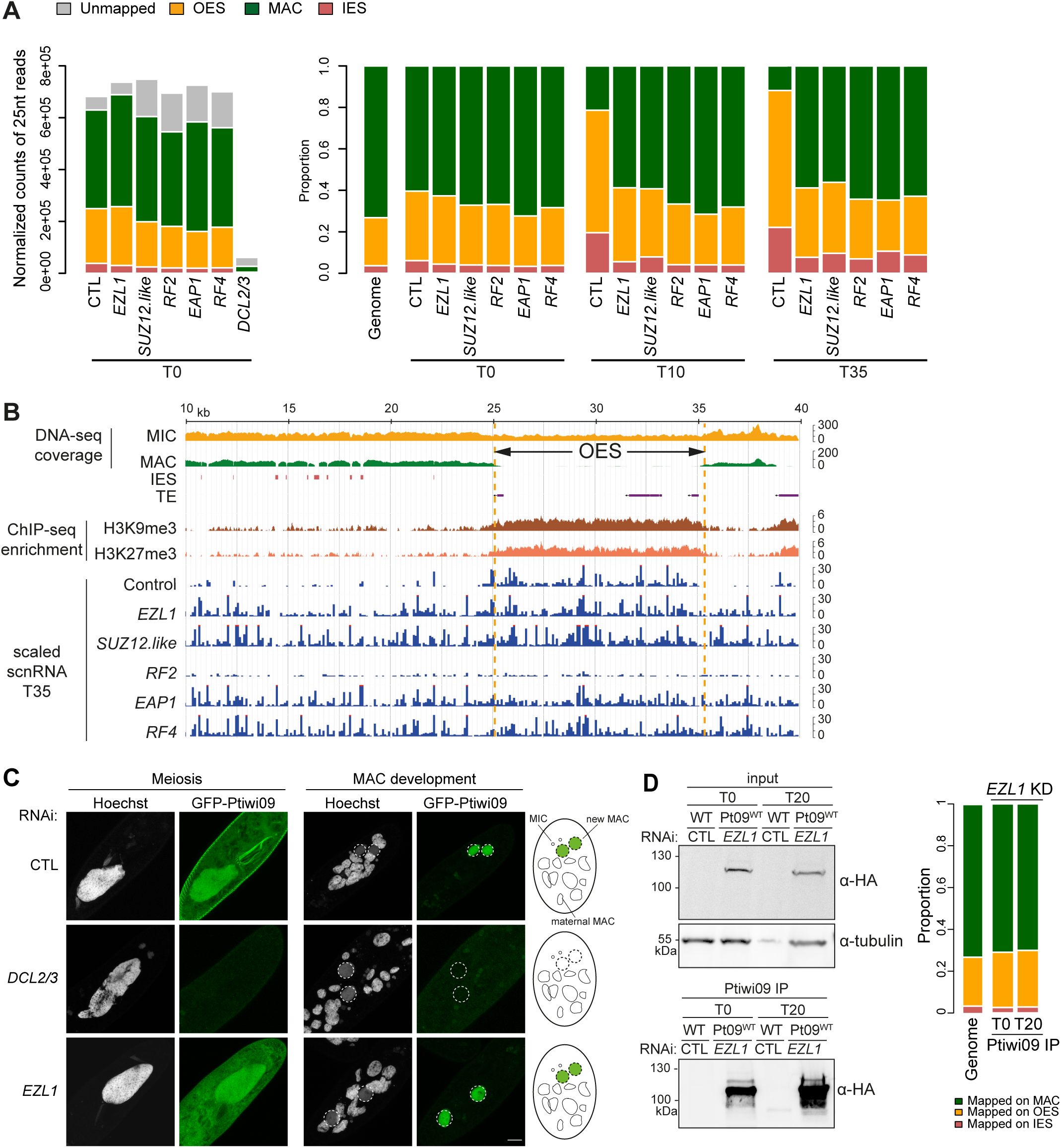
PRC2-Ezl1 core complex and its associated cofactors are required for scnRNA selection. A. Left: Normalized counts of 25nt-long scnRNAs at T0 in ICL7 (CTL), EZL1, SUZ12.like, RF2, EAP1, RF4 and DCL2/3 RNAi conditions. Bar plots show the reads that are unmapped or match the MAC genome, IESs and OES reference sequences. Right: Proportion of 25nt-long scnRNAs that match the MAC genome, IESs and OES reference sequences at different time points of the sexual cycle in control (CTL), EZL1, SUZ12.like, RF2, EAP1, and RF4 RNAi conditions. Genome: proportion of each category (MAC, OES, IES) in the MIC genome. B. Representative genomic region (same as in Figure 2B) depicting 25-nt scnRNAs at T35 in control (CTL), EZL1, SUZ12.like, RF2, EAP1, and RF4 RNAi. DNA-seq coverage (bin=1) for the MIC and the MAC DNA samples on the MIC genome (NODE_1273_length_39901_cov_28.471216 between 10 and 40 kb), and IES and TE annotations are indicated. ChIP-seq enrichment (deeptools bamCompare bin=5) of H3K9me3 and H3K27me3 in PGM RNAi conditions (data from (19)) is displayed. Scaled scnRNA coverages (bin=100) were calculated on equivalent read datasets (15e6 sRNA between 20 and 30-nt long) for all samples except RF2, for which all available reads (3.2e6) were considered. C. GFP-Ptiwi09 localization in CTL, DCL2/3 and EZL1 RNAi conditions in meiosis and during MAC development. Representative confocal images are displayed. Overlay of Z-projections are presented. Dashed white circles indicate the new developing MACs. Scale bar is 10 μm. Note that, in contrast to control cells, basal bodies on the cell cortex are not decorated with GFP fluorescence in Ezl1-depleted cells. D. Sequencing analysis of scnRNA populations after Ptiwi09 immunoprecipitation in EZL1 KD. Left: Western blot analysis of whole cell extracts at T0 and T20 from cells expressing the 3XFLAG-HA-Ptiwi09 protein (Pt09WT) or wild-type cells (WT), before (input) and after FLAG immunoprecipitation (Ptiwi09 IP) upon EZL1 RNAi or no RNAi (CTL). Anti-HA antibodies are used to detect Ptiwi09 and alpha-tubulin for normalization. Right: Proportion of 25nt-long scnRNAs that match the MAC genome, IES and OES reference sequences after Ptiwi09 IP at T0 and T20 in EZL1 RNAi. Genome: proportion of each category (MAC, OES, IES) in the MIC genome.

Importantly, because we still detect maternal MAC ncRNA production in the *EZL1* knockdown experiments, the lack of scnRNA selection cannot be explained by a lack of maternal MAC ncRNA transcription (Supplementary Figure S5). To determine whether impairment of scnRNA selection could be explained by defects in the localization of scnRNAs, we analyzed the localization of the scnRNA-binding protein Ptiwi09 with a *GFP-PTIWI09* transgene. In control cells, the GFP-Ptiwi09 fusion protein was detected in the cytoplasm and in the maternal MAC during MIC meiosis then localized exclusively in the developing MAC, as expected (31)(Figure 4C). In contrast, in Dcl2/3-depleted cells, in which scnRNA biogenesis is abrogated, the GFP-Ptiwi09 fusion protein was no longer detected during MIC meiosis and MAC development (Figure 4C). This observation is consistent with unloaded Argonaute proteins being generally unstable (52, 53). Thus, GFP-Ptiwi09 localization likely reflects scnRNA localization. In Ezl1-depleted cells, where scnRNA selection is defective, GFP-Ptiwi09 still localized in the cytoplasm and in the maternal MAC during MIC meiosis, and then in the new developing MACs, as in control conditions (Figure 4C). We conclude that the lack of MIC-specific scnRNA selection in the absence of Ezl1 does not appear to be associated with the mis-localization of scnRNAs.

To assess whether unselected scnRNAs still bind to Ptiwi09, we performed Ptiwi09 immunoprecipitation in *EZL1* RNAi conditions (Figure 4D) followed by small RNA sequencing, when Ptiwi09 is present in the maternal MAC (T0) and when it is in the new developing MAC (T20). 25 nt-scnRNAs are present at both time points (Supplementary Figure S5) and the same proportions of MIC-and MAC scnRNAs are detected in the IP at both time points (Figure 4D). This indicates that the non-selected scnRNAs that accumulate in Ezl1-depleted cells are bound to Ptiwi09.

### H3K27me3 in the maternal MAC is dispensable for scnRNA selection

PRC2-Ezl1 catalyzes H3K27me3 in the maternal MAC (21, 22). To directly assess whether H3K27me3 is required for scnRNA selection, we sought to identify an experimental condition where H3K27me3 no longer accumulates in the maternal MAC. We designed a mutant H3 transgene carrying a lysine to methionine substitution in position 27 (H3K27M), a mutation described to cause a dominant negative effect preventing H3K27me3 accumulation at the endogenous H3 by inhibiting the enzymatic activity of PRC2 and resulting in genome-wide depletion of H3K27me3 (54, 55).

Five H3 proteins are encoded in the *Paramecium tetraurelia* MAC genome. H3P1 and H3P3 are H3.1 histones and are constitutively expressed during the life cycle (56). We transformed *Paramecium* cells with histone *H3P1* or *H3P3* transgenes bearing K27M mutations and induced sexual events. Transformation with *H3P1_K27M_* or *H3P3_K27M_* transgenes abolished H3K27me3 accumulation in the maternal MAC without affecting H3K27me3 deposition in the MIC during meiosis I or in new developing MACs based on immunofluorescence experiments (Figure 5A, Supplementary Figure S6). This does not appear to be due to a mislocalization of Ezl1 because in cells co-transformed with *H3P3_K27M_* and *FLAG-EZL1* transgenes, the fusion protein FLAG-Ezl1 localized in the MIC and the maternal MAC during meiosis I then in new developing MACs (Supplementary Figure S6), as was previously reported for a *GFP-EZL1* transgene (22). The *H3P1_K27M_* or *H3P3_K27M_* transgenes had no major effect on the accumulation of H3K9me3 in the new MAC (Supplementary Figure S6). Testing elimination of some IESs did not reveal any DNA elimination defect (Supplementary Figure S6) and the survival of the sexual progeny was not affected (Figure 5B), indicating that deposition of H3K27me3 in the maternal MAC did not appear to be required for IES elimination and viability of the sexual progeny. We sequenced the small RNA populations at two time points in *H3P1_K27M_* transformed cells (Supplementary Figure S6). The scnRNAs were produced normally at early developmental stages (T0) and were enriched in MIC-specific sequences at later stages (T20) (Figure 5C and Supplementary Figure S6). Thus, the scnRNA selection process does not appear to require H3K27 trimethylation in the maternal MAC. Consistent with this idea, H3K27me3 deposition is not affected in the maternal MAC of Eap1-depleted cells (19), yet scnRNA selection is impaired (Figure 4A-B), suggesting that PRC2-Ezl1 integrity is critical for scnRNA selection for other reasons.

**Figure 5.**
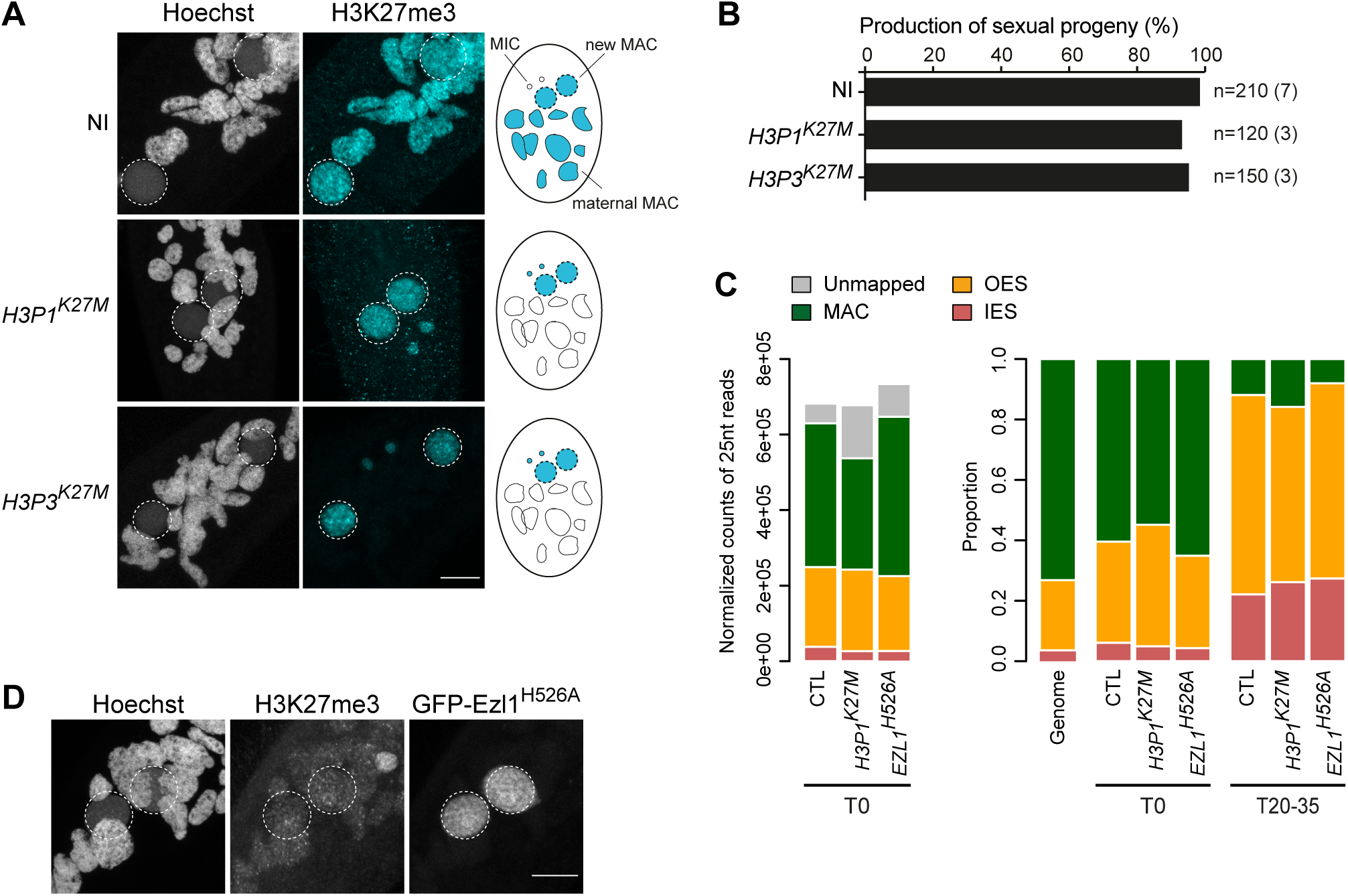
Ezl1 histone methyltransferase activity is not required for scnRNA selection. A. H3K27me3 immunostaining of non-injected cells (NI), and of cells transformed with H3P1K27M or H3P3K27M during MAC development. Representative confocal images are displayed. Overlay of Z-projections are presented. Dashed white circles indicate the new developing MACs. Note the abnormal accumulation of H3K27me3 in the new MICs in cells transformed by H3P1K27M or H3P3K27M, or with GFP-EZL1H526A mutant transgene (panel D). The same abnormal localization was observed in RNAi-mediated Ezl1-depleted cells (21). Scale bar is 10 μm. B. Production of sexual progeny in cells transformed with H3P1K27M or H3P3K27M. The total number of analyzed cells and the number of independent experiments (in parenthesis) are indicated. C. Left: Normalized counts of 25-nt scnRNAs at T0 in control (CTL) RNAi, in cells transformed with H3P1K27M, and in cells transformed with GFP-EZL1H526A upon endogenous EZL1 RNAi. Bar plots show the reads that are unmapped or match the MAC genome, IES and OES reference sequences. Right: Proportion of 25-nt scnRNAs that match the MAC genome, IES and OES reference sequences at T0 and T20-35 of the sexual cycle. Genome: proportion of each category (MAC, OES, IES) in the MIC genome. E. H3K27me3 immunostaining and GFP-EZL1H526A localization during new MAC development in cells transformed with GFP-EZL1H526A upon endogenous EZL1 RNAi. Representative confocal images are displayed. Overlay of Z-projections of magnified views of Hoechst staining, H3K27me3-specific antibodies and GFP signal are presented. Dashed white circles indicate the new developing MACs. Note the abnormal accumulation of H3K27me3 in the new MICs in cells transformed with the GFP-EZL1H526A mutant transgene. Scale bar is 10 μm.

### Ezl1 histone methyltransferase activity is not required for scnRNA selection

Our data indicate that PRC2-Ezl1, but not its signature H3K27me3 in the maternal MAC, is necessary for scnRNA selection. We hypothesized that PRC2-Ezl1 may function in scnRNA selection by methylating substrates other than H3K27. To test this possibility, we used a catalytic mutant of Ezl1. The H526A mutation in the SET domain of Ezl1 was previously shown to abolish Ezl1 catalytic activity *in vitro* and *in vivo* (21). In cells expressing the RNAi-resistant *GFP-EZL1_H526A_* mutant transgene and silenced for the endogenous *EZL1* gene, H3K27me3 no longer accumulated in the maternal MAC nor in the new developing MACs, even though the mutant Ezl1 protein still correctly localized (Figure 5D, Supplementary Figure S6). In this context, we performed *s*mall RNA sequencing. Contrary to our expectations, scnRNAs were produced normally at the beginning of the sexual cycle (T0) and MIC-specific scnRNAs were correctly selected at later stages (T35) (Figure 5C and Supplementary Figure S6). Thus, we conclude that the function of PRC2-Ezl1 in scnRNA selection is independent of its histone methyltransferase activity.

### The PRC2 cofactor Rf4 is required for the interaction between Ptiwi09 and Gtsf1 in the maternal MAC

Previous work showed that the Ptiwi09 protein interacts in the maternal MAC with the tandem zinc finger protein Gtsf1, which is essential for scnRNA selection, and PRC2 (24, 25). We therefore reasoned that the interaction between Ptiwi09 and Gtsf1 might be mediated by PRC2. To test this hypothesis, we compared the proteins interacting with Gtsf1 in control conditions with those found in cells depleted for Rf4, the PRC2 cofactor known to bridge PRC2 to Ptiwi09 in the new developing MAC (19). The localization in the maternal MAC of the 3xFLAG-HA-Gtsf1 fusion protein in control and Rf4-depleted cells was confirmed by immunofluorescence (Figure 6A). The Gtsf1 nuclear signal appears lower in *RF4* RNAi compared to control RNAi, based on immunofluorescence and western blot analysis of nuclear extracts (Figure 6A, Supplementary Figure S7). Yet, Rf4 is not required for Gtsf1 expression, given the similar levels of the Gtsf1 fusion protein in whole cells extracts from control and Rf4-depleted cells (Supplementary Figure S7). Immunoprecipitation (IP) of the nuclear Gtsf1 fusion protein, in 3 replicates for the control and Rf4-depleted cells, respectively (Materials and Methods), was followed by quantitative label-free mass spectrometry (Figure 6B). Statistical analyses revealed 161 differential proteins (out of 2613 identified proteins) in Rf4-depleted cells compared with control (fold change ≥ 2; p value ≤ 0.05; unique peptide ≥ 2) (Figure 6B; Supplementary Figure S7). We recovered Gtsf1 in both conditions, as expected (Figure 6B). Among the 64 proteins that were significantly depleted upon *RF4* RNAi, not only did we find Rf4 as expected, but we also found PRC2 components (Ezl1, Eed, Suz12-like) and cofactors (Rf2 and Eap1), Ptiwi09 and its interacting proteins (Ptiwi01, Ptiwi03 and Ema1a) (Figure 6B) (19). This result is in agreement with a previous study reporting that PRC2 composition at early stages of autogamy includes Ezl1, Caf1, Eed, Suz12-like, Rf2 and Rf4 (20). Therefore, we conclude that depletion of the PRC2-Ezl1 cofactor Rf4 disrupts the interaction between Gtsf1, Ptiwi09 and PRC2.

**Figure 6.**
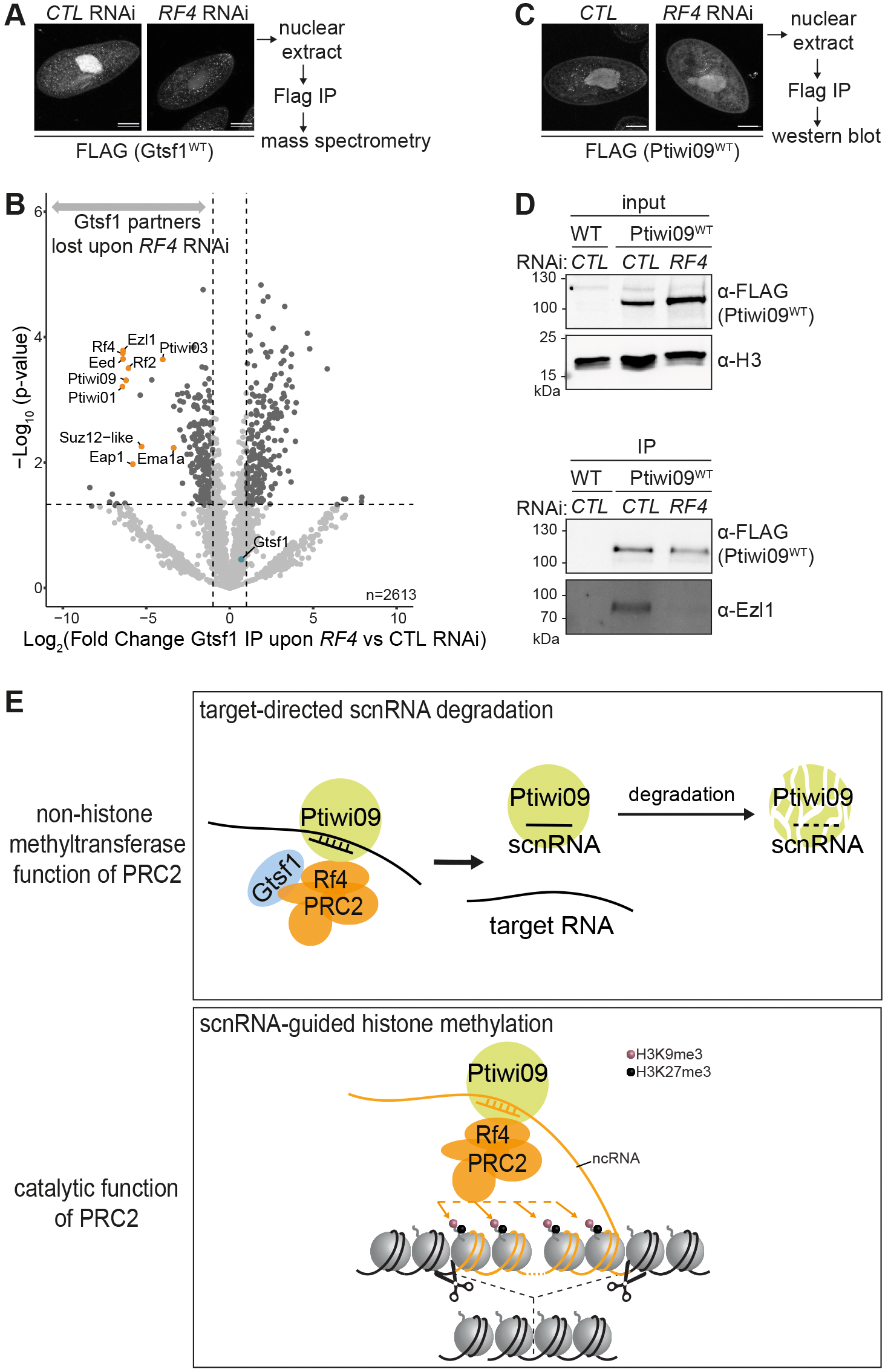
The PRC2 cofactor Rf4 is required for the interaction between Ptiwi09 and Gtsf1 in the maternal MAC. **A**. Gtsf1 immunoprecipitation experiments upon /CL7 (CTL) or RF4 RNAi. Anti-FLAG immunostaining of cells transformed with a 3XFLAG-HA-GTSF1 transgene are shown, at the same time point as the preparation of nuclear extracts for FLAG immunoprecipitation (TO) followed by quantitative mass spectrometry. Scale bar, 10 µm. B. Volcano plot of the quantitative label-free mass-spectrometry analysis of 3xFLAG-HA-Gtsf1 affinity purification upon RF4 vs ICU (CTL) RNAi. 3 replicates were analyzed for each condition. Significantly enriched proteins in Gtsf1 IP upon RF4 RNAi over Gtsf1 IP upon CTL RNAi are shown in dark grey (statistical I-test), Fold Change<!: 2, and p-value s 0.05, only proteins identified with more than 2 unique peptides are considered. Gtsf1 is highlighted in blue. PRC2 components and Ptiwi09-associated proteins are highlighted in yellow. C. Ptiwi09 immunoprecipitation experiments upon CTL (no RNAi) or RF4 RNAi. Anti-FLAG immunostaining of cells transformed with a 3XFLAG-HA-PTIW/09 transgene are shown, at the same time point as the preparation of nuclear extracts for FLAG immunoprecipitation (TO). Scale bar, 10 µm. D. Western blot analysis of nuclear extracts from cells expressing the 3XFLAG-HA-Ptiwi09 protein (Ptiwi09wT) or 3XFLAG-HA (CTL) before (input) or after immunoprecipitation (IP). Anti-FLAG antibodies are used to detect Ptiwi09, anti-Ezl1 to detect Ezl1, and anti-histone H3 antibodies for normalization. E. Working model. The presence of PRC2 core subunits and cofactors in the maternal MAC but not the deposition of H3K27me3 appears to be required for the degradation of MAC-scnRNAs. The non-histone methyltransferase function of PRC2-Ezl1 in the degradation of MAC-scnRNAs may contribute to the stabilization of the interaction between MAC-scnRNA/Ptiwi09 complexes and nascent non-coding RNAs. Through an architectural role, PRC2-Ezl1 would bring together Ptiwi09 and Gtsf1 through the Rf4 protein, leading to Ptiwi09 ubiquitylation and degradation via the proteasome.

To assess whether the interaction between Gtsf1 and PRC2-Ezl1 might involve RNA, we examined Gtsf1 IP after control and RNase I treatment, in two independent biological replicates (Supplementary Figure S7). We detected Ezl1 in the Gtsf1 IP but not in the control IP, as expected. After treatment with RNase I, Ezl1 was still found in the Gtsf1 IP, consistent with a RNA-independent interaction between Gtsf1 and PRC2-Ezl1. This observation is in agreement with previous work showing that the interaction between Gtsf1 and Ptiwi09 is RNA-independent (24).

Given that Rf4 impairs Gtsf1 localization in the maternal MAC, we examined the localization and levels of Ptiwi09 fusion protein in control and Rf4-depleted cells. Western blot analysis of whole cells extracts and immunostaining indicated that *RF4* RNAi has no effect on Ptiwi09 (Supplementary Figure S7).

To assess whether Rf4 is required for the interaction between Ptiwi09 and PRC2-Ezl1 in the maternal MAC, we performed Ptiwi09 immunoprecipitation in control, and *RF4* RNAi conditions and examined the presence of Ezl1 by western blot. Cells expressing the 3xFLAG-HA-Ptiwi09 fusion protein in the maternal MAC were used for the immunoprecipitation (IP) of the FLAG tag from nuclear extracts (Figure 6C; Supplementary Figure S7). Western blot analysis indicated that Ezl1 is found in the control Ptiwi09 IP, as expected, while it is not detected in Rf4-depleted cells (Figure 6D). Consistently, we found that Ezl1 is no longer detected in the maternal MAC upon *RF4* RNAi (Supplementary Figure S7). Thus, the interaction between Ptiwi09 and Ezl1 is mediated, directly or indirectly, by the PRC2 cofactor Rf4 in the maternal MAC, as it is the case in the new developing MAC (19).

## Discussion

This study provides a comprehensive description of scnRNA populations during the sexual process of autogamy in *Paramecium tetraurelia* (Figure 2). Using cells arrested in meiosis I, we demonstrate that scnRNAs begin accumulating prior to or during early meiosis I. Covering the entire sexual cycle of autogamy with sRNA sequencing at 11 time points, we determined, at an unprecedented resolution, that scnRNAs are produced from both strands for the whole MIC genome in *Paramecium tetraurelia*, with no bias in scnRNA coverage between IESs and MAC-destined sequences. The mechanism underlying this genome-wide transcription is being thoroughly studied but remains elusive (57, 58).

After scnRNAs are produced from the entire MIC genome, genome-wide degradation of MAC-scnRNAs occurs; the earliest time at which degradation is detected corresponds to the onset of DNA elimination (Figure 2). As a result of this specific degradation, the subpopulation of scnRNAs corresponding to MIC-specific sequences is selected and persists until the end of MAC development. The dynamics of scnRNAs we observed during *Paramecium* sexual events contrasts with that described in another ciliate, *Tetrahymena thermophila* (5). Indeed, during *Tetrahymena* conjugation, scnRNAs are preferentially produced from eliminated sequences, resulting in a biased production of scnRNAs from MIC-specific sequences. In addition, there is a rather limited selection process. In contrast, the massive degradation of MAC-scnRNAs (> 70 Mb of complexity) makes *Paramecium* a very good model to decipher the mechanisms involved in scnRNA selection.

Here, we showed that RNAi-mediated knockdown of genes encoding core components of the PRC2-Ezl1 core complex (Ezl1 and Suz12.like) and its associated cofactors (Rf2, Rf4 and Eap1) prevented scnRNA selection (Figure 4). Thus, the PRC2-Ezl1 complex is involved in MAC-scnRNA degradation, in agreement with previous studies (20, 51). Ezl1 has been shown to transiently localize in the maternal MAC, where the selection process likely occurs, before it is detected in the new developing MACs (22). Likewise, Ezl1-interacting partners are also transiently localized in the maternal MAC before they localize in the new MAC (19, 20). (20) report that the PRC2 complex exhibits a similar composition in the maternal MAC and the new MAC. Consistent with these observations, pull down of the Gtsf1 protein localized in the maternal MAC (Figure 6) identified known core PRC2 components and cofactors, including Eap1 which was not detected in (20). We thus conclude that PRC2-Ezl1 function in scnRNA selection requires the integrity of the complex (Figure 4), its interaction with Ptiwi09 via Rf4 (Figure 6), and the accessory factor Eap1 (Figure 4), whose exact role remains to be determined.

Monitoring the localization of the scnRNA-binding protein Ptiwi09 indicated that the scnRNA-Ptiwi09 complexes are correctly localized, despite the lack of selection in PRC2-Ezl1 depleted cells (Figure 4). Consistently, Ptiwi09 proteins present in the new MAC are loaded with both MAC-and MIC-scnRNAs in Ezl1-depleted cells (Figure 4). Thus, in absence of selection, both MAC-and MIC-scnRNAs are present in the new MAC, a situation that is radically different from wild-type cells, where only MIC-scnRNAs are present in the new MAC and guide histone modifications at target DNA sequences (19). We conclude that the mere presence of scnRNA-Ptiwi09 complexes in the new MAC does not appear to be sufficient to trigger DNA elimination. This confirms that the availability of PRC2-Ezl1 components in the new MAC is also critical for DNA elimination to occur, as implied by recent work from us and others (19, 20).

We showed that the specific abrogation of H3K27me3 in the maternal MAC, the signature of PRC2-Ezl1, had no impact on scnRNA selection, demonstrating that this mark does not play a role in this process (Figure 5). To suppress H3K27me3, we expressed dominant negative forms of the H3.1 protein in which lysine 27 is replaced by a methionine (H3P1_K27M_ and H3P3_K27M_) (Figure 5). These mutant proteins prevented the accumulation of H3K27me3 in the maternal MAC, but not in the new MAC, even though the mutated H3 and the PRC2 catalytic subunit Ezl1 were detected in both nuclei (Supplementary Figure S6). The reason why the dominant negative effect of the H3.1 mutant protein is restricted to the maternal MAC is not known. The molecular mechanism by which H3K27M reduces H3K27me2/3 in cells remains a matter of debate more generally (59, 60). Post-translational modifications of the same H3 tail, both proximal and distal to K27M, can greatly diminish the inhibition of PRC2 (61). Furthermore, the H3K27M-to-PRC2 stoichiometric ratio is believed to dictate the degree of PRC2 inhibition (62). Thus, we speculate that H3.1 mutant proteins are unable to exert a dominant negative effect in the developing MACs because (i) H3P1 and H3P3 are not the substrates of PRC2-Ezl1 in the developing MACs; other H3 proteins are expressed during MAC development (56); (ii) H3P1 and H3P3 in the developing MACs bear post-translational modifications that impede the dominant negative effect of the H3K27M mutation; or (iii) the amounts of PRC2 relative to H3K27M in the developing MACs are too high to be sensitive to H3K27M-mediated inhibition of PRC2 activity.

We did not observe any apparent consequence of the lack of H3K27me3 in the maternal MAC on DNA elimination and sexual progeny production. Therefore, we conclude that H3K27me3 deposition in the maternal MAC is not required for DNA elimination or sexual progeny production. At this point, the role of H3K27me3 in the maternal MAC remains elusive. If H3K27me3 controls gene expression, it might not involve the expression of genes necessary for the completion of sexual events.

Our findings prompted us to investigate whether PRC2-Ezl1 might function through a different mechanism, independent of H3K27 methylation. Like many histone-modifying enzymes, the mammalian PRC2 has non-histone substrates, including the nuclear receptor Retinoid-Related Orphan Receptor Alpha (63), the transcription factor GATA4 (64), the facultative PRC2 cofactors JARID2 (65) and PALI1 (66), and the RNAPII transcription elongation factor Elongin A (67). To assess whether PRC2-Ezl1 function in the scnRNA selection process may be decoupled from its methyltransferase activity, we tested a catalytic dead mutant of Ezl1, carrying a histidine-to-alanine substitution in its catalytic SET domain (H526A) (21). We showed that scnRNA selection occurred normally in the *EZL1_H526A_* catalytic mutant, demonstrating that PRC2-Ezl1 has a function independent of its histone methyltransferase activity (Figure 5). One example of a non-catalytic function of the mammalian PRC2-Ezh1 was previously shown in an experimental setup where the artificial recruitment of a catalytically inactive Ezh1 led to transcriptional repression of a reporter via chromatin compaction (68).

The scanning model proposes that the discrimination between MAC-and MIC-scnRNA/Ptiwi09 complexes results from the pairing of scnRNAs with complementary nascent non-coding transcripts produced by the maternal MAC. Only MAC-scnRNAs are degraded. One potential mechanism involves the post-transcriptional modification 2’-O-methylation at the 3’ terminus by the methyl transferase HEN1. *Tetrahymena* scnRNAs, like piRNAs in metazoans, are 2’-O-methylated at their 3’ ends (69, 70). Hen1p-mediated 2’-O-methylation was reported to stabilize scnRNAs. Loss of Hen1p causes a reduction in the abundance and length of scnRNAs, defects in programmed genome elimination, and low production of sexual progeny (70). One could imagine that the scanning process leads to the differential 2’-O-methylation of MAC-and MIC-scnRNAs: MIC-scnRNAs would be 2’-O-methylated, which would protect them from degradation, while MAC-scnRNAs would not. The re-analysis of published data (38) contradicts this hypothesis, however, because *HEN1* KD during *Paramecium* autogamy did not impair scnRNA selection (Supplementary Figure S8). Thus, 2’-O-methylation of scnRNAs does not appear to be involved in the selection process.

Only MAC-scnRNAs display extensive complementarity with the non-coding RNA targets. We propose that the pairing between MAC-scnRNAs and MAC non-coding RNA leads to the degradation of the Ptiwi09 protein and its bound scnRNAs. In agreement with this model, recent work provided evidence that Ptiwi09 is ubiquitylated in a Gtsf1-dependent manner and Ptiwi09 degradation involves the proteasome (24). In essence, the proposed mechanism is similar to microRNA target-directed degradation, in which pairing of the miRNA to the cognate RNA leads to ubiquitination and proteasomal degradation of Ago and of the miRNA (71, 72). Based on our findings, we propose that the non-histone methyltransferase function of PRC2-Ezl1 in the degradation of MAC-scnRNAs contributes to the stabilization of the interaction between MAC-scnRNA/Ptiwi09 complexes and nascent non-coding RNAs.

What could be the role of PRC2 in scnRNA selection? We show that the PRC2 cofactor Rf4 interacts, directly or indirectly, with Gtsf1 and Ptiwi09 (Figure 6B), and that Rf4 controls Gtsf1 nuclear localization (Figure 6A, Supplementary Figure S7), but not that of Ptiwi09 (Supplementary Figure S7). We envision two non-exclusive hypotheses for the role of PRC2 that takes into account its interaction with Gtsf1 and Ptiwi09. In the first hypothesis, PRC2 may bring together Ptiwi09 and Gtsf1, suggestive of an architectural role for PRC2-Ezl1. The physical interaction mediated by the PRC2 cofactor Rf4 might be necessary for the nuclear anchoring of Gtsf1 in the maternal MAC. The impaired nuclear localization of Gtsf1 in Rf4-depleted cells would thus be a consequence of the loss of interaction between Rf4 and Gtsf1 (Figure 6B). If the physical interaction between Ptiwi09 and Gtsf1 requires Rf4, it raises the possibility that Ptiwi09 may not interact directly with Gtsf1, unlike the direct binding of PIWI and GTSF1 observed in *Drosophila* and mice (73, 74). In the second hypothesis, the PRC2 cofactor Rf4 may be important to trigger a conformational change or a post-translational modification (other than lysine methylation) required for Gtsf1-Ptiwi09 interaction. Loss of this interaction, in the absence of Rf4, would impair Gtsf1 anchoring in the nucleus. Whether Rf4, Gtsf1 and Ptiwi09 interact directly or indirectly, and how these proteins interact with each other will require further investigation.

In addition to its catalytic role in the new MAC, where it works in concert with Ptiwi09 to deposit histone marks on transposable elements (19), our study reveals that PRC2 performs a distinct, histone methyltransferase-independent function in the maternal MAC, where, in association with Ptiwi09 and Gtsf1, it participates in scnRNA degradation (Figure 6E). Future work will be needed to elucidate the mechanisms by which PRC2-Ezl1 exerts its function independently of its histone methyltransferase activity and whether they are analogous to those involved in microRNA target-directed degradation.

## ACKNOWLEDGEMENTS

We wish to thank the members of the Duharcourt lab for fruitful discussions, and Sébastien Léon for his insightful advice. We thank Daniel Holoch and Eric Meyer for comments on the manuscript. We acknowledge the ImagoSeine facility, member of the FranceBioImaging infrastructure supported by the ANR-10-INSB-04, and the sequencing and bioinformatics expertise of the I2BC High-throughput sequencing facility, supported by France Génomique (funded by the French National Program “Investissement d’Avenir” ANR-10-INBS-09).

## AUTHOR CONTRIBUTIONS

CMP, OC, MG conducted most experiments with the help of AH for cloning and silencing experiments; AV with the help of MLA performed sRNA time course experiments; CMP and AF analyzed the Ezl1 catalytic mutant; OA designed and performed the bioinformatic analyses of NGS data; GC performed the mass spectrometry analyses; CMP, OC, MG, SD designed the experiments, CMP, OC, MG prepared the figures and SD wrote the paper with input from all co-authors. SD supervised the project.

## REFERENCES

1. Han, J. and Mendell, J.T. (2023) MicroRNA turnover: a tale of tailing, trimming, and targets. Trends Biochem. Sci., 48, 26–39.

2. Lepere, G., Nowacki, M., Serrano, V., Gout, J.F., Guglielmi, G., Duharcourt, S. and Meyer, E. (2009) Silencing-associated and meiosis-specific small RNA pathways in Paramecium tetraurelia. Nucleic Acids Res, 37, 903–15.

3. Mochizuki, K., Fine, N.A., Fujisawa, T. and Gorovsky, M.A. (2002) Analysis of a piwi-related gene implicates small RNAs in genome rearrangement in tetrahymena. Cell, 110, 689–99.

4. Sandoval, P.Y., Swart, E.C., Arambasic, M. and Nowacki, M. (2014) Functional Diversification of Dicer-like Proteins and Small RNAs Required for Genome Sculpting. Dev. Cell, 28, 174–188.

5. Schoeberl, U.E., Kurth, H.M., Noto, T. and Mochizuki, K. (2012) Biased transcription and selective degradation of small RNAs shape the pattern of DNA elimination in Tetrahymena. Genes Dev., 26, 1729–1742.

6. Singh, D.P., Saudemont, B., Guglielmi, G., Arnaiz, O., Goût, J.-F., Prajer, M., Potekhin, A., Przybòs, E., Aubusson-Fleury, A., Bhullar, S., et al. (2014) Genome-defence small RNAs exapted for epigenetic mating-type inheritance. Nature, 509, 447–452.

7. Betermier, M. and Duharcourt, S. (2014) Programmed Rearrangement in Ciliates: Paramecium. Microbiol. Spectr., 2.

8. Aury, J.M., Jaillon, O., Duret, L., Noel, B., Jubin, C., Porcel, B.M., Segurens, B., Daubin, V., Anthouard, V., Aiach, N., et al. (2006) Global trends of whole-genome duplications revealed by the ciliate Paramecium tetraurelia. Nature, 444, 171–8.

9. Guérin, F., Arnaiz, O., Boggetto, N., Denby Wilkes, C., Meyer, E., Sperling, L. and Duharcourt, S. (2017) Flow cytometry sorting of nuclei enables the first global characterization of Paramecium germline DNA and transposable elements. BMC Genomics, 18, 327.

10. Sellis, D., Guérin, F., Arnaiz, O., Pett, W., Lerat, E., Boggetto, N., Krenek, S., Berendonk, T., Couloux, A., Aury, J.-M., et al. (2021) Massive colonization of protein-coding exons by selfish genetic elements in Paramecium germline genomes. PLOS Biol., 19, e3001309.

11. Swart, E.C., Denby Wilkes, C., Sandoval, P.Y., Hoehener, C., Singh, A., Furrer, D.I., Arambasic, M., Ignarski, M. and Nowacki, M. (2017) Identification and analysis of functional associations among natural eukaryotic genome editing components. F1000Research, 6, 1374.

12. Duharcourt, S., Keller, A.M. and Meyer, E. (1998) Homology-dependent maternal inhibition of developmental excision of internal eliminated sequences in Paramecium tetraurelia. Mol Cell Biol, 18, 7075–85.

13. Duharcourt, S., Butler, A. and Meyer, E. (1995) Epigenetic self-regulation of developmental excision of an internal eliminated sequence on Paramecium tetraurelia. Genes Dev, 9, 2065–77.

14. Furrer, D.I., Swart, E.C., Kraft, M.F., Sandoval, P.Y. and Nowacki, M. (2017) Two Sets of Piwi Proteins Are Involved in Distinct sRNA Pathways Leading to Elimination of Germline-Specific DNA. Cell Rep., 20, 505–520.

15. Aronica, L., Bednenko, J., Noto, T., DeSouza, L.V., Siu, K.W., Loidl, J., Pearlman, R.E., Gorovsky, M.A. and Mochizuki, K. (2008) Study of an RNA helicase implicates small RNA-noncoding RNA interactions in programmed DNA elimination in Tetrahymena. Genes Dev, 22, 2228–41.

16. Lepere, G., Betermier, M., Meyer, E. and Duharcourt, S. (2008) Maternal noncoding transcripts antagonize the targeting of DNA elimination by scanRNAs in Paramecium tetraurelia. Genes Dev, 22, 1501–12.

17. Martienssen, R. and Moazed, D. (2015) RNAi and Heterochromatin Assembly. Cold Spring Harb. Perspect. Biol., 7, a019323.

18. Maliszewska-Olejniczak, K., Gruchota, J., Gromadka, R., Denby Wilkes, C., Arnaiz, O., Mathy, N., Duharcourt, S., Bétermier, M. and Nowak, J.K. (2015) TFIIS-Dependent Non-coding Transcription Regulates Developmental Genome Rearrangements. PLoS Genet., 11, e1005383.

19. Miró-Pina, C., Charmant, O., Kawaguchi, T., Holoch, D., Michaud, A., Cohen, I., Humbert, A., Jaszczyszyn, Y., Chevreux, G., Maestro, L.D., et al. (2022) Paramecium Polycomb repressive complex 2 physically interacts with the small RNA-binding PIWI protein to repress transposable elements. Dev. Cell, 57, 1037–1052.e8.

20. Wang, C., Solberg, T., Maurer-Alcalá, X.X., Swart, E.C., Gao, F. and Nowacki, M. (2022) A small RNA-guided PRC2 complex eliminates DNA as an extreme form of transposon silencing. Cell Rep., 40.

21. Frapporti, A., Miró Pina, C., Arnaiz, O., Holoch, D., Kawaguchi, T., Humbert, A., Eleftheriou, E., Lombard, B., Loew, D., Sperling, L., et al. (2019) The Polycomb protein Ezl1 mediates H3K9 and H3K27 methylation to repress transposable elements in Paramecium. Nat. Commun., 10, 2710.

22. Lhuillier-Akakpo, M., Frapporti, A., Denby Wilkes, C., Matelot, M., Vervoort, M., Sperling, L. and Duharcourt, S. (2014) Local effect of enhancer of zeste-like reveals cooperation of epigenetic and cis-acting determinants for zygotic genome rearrangements. PLoS Genet., 10, e1004665.

23. Allen, S.E., Hug, I., Pabian, S., Rzeszutek, I., Hoehener, C. and Nowacki, M. (2017) Circular Concatemers of Ultra-Short DNA Segments Produce Regulatory RNAs. Cell, 168, 990.

24. Charmant, O., Gruchota, J., Arnaiz, O., Nowak, K.P., Moisan, N., Zangarelli, C., Bétermier, M., Anielska-Mazur, A., Legros, V., Chevreux, G., et al. (2024) The PIWI-interacting protein Gtsf1 controls the selective degradation of small RNAs in Paramecium. Nucleic Acids Res., 10.1093/nar/gkae1055.

25. Wang, C., Lyv, L., Solberg, T., Zhang, H., Wen, Z. and Gao, F. (2024) GTSF1 is required for transposon silencing in the unicellular eukaryote Paramecium tetraurelia. Nucleic Acids Res., 10.1093/nar/gkae925.

26. Marker, S., Carradec, Q., Tanty, V., Arnaiz, O. and Meyer, E. (2014) A forward genetic screen reveals essential and non-essential RNAi factors in Paramecium tetraurelia. Nucleic Acids Res., 42, 7268– 7280.

27. Beisson, J., Betermier, M., Bre, M.H., Cohen, J., Duharcourt, S., Duret, L., Kung, C., Malinsky, S., Meyer, E., Preer, J.R., et al. (2010) Maintaining clonal Paramecium tetraurelia cell lines of controlled age through daily reisolation. Cold Spring Harb Protoc, 2010, pdb prot5361.

28. Beisson, J., Betermier, M., Bre, M.H., Cohen, J., Duharcourt, S., Duret, L., Kung, C., Malinsky, S., Meyer, E., Preer, J.R., et al. (2010) Mass culture of Paramecium tetraurelia. Cold Spring Harb Protoc, 2010, pdb prot5362.

29. Garnier, O., Serrano, V., Duharcourt, S. and Meyer, E. (2004) RNA-mediated programming of developmental genome rearrangements in Paramecium tetraurelia. Mol Cell Biol, 24, 7370–9.

30. Baudry, C., Malinsky, S., Restituito, M., Kapusta, A., Rosa, S., Meyer, E. and Betermier, M. (2009) PiggyMac, a domesticated piggyBac transposase involved in programmed genome rearrangements in the ciliate Paramecium tetraurelia. Genes Dev, 23, 2478–83.

31. Bouhouche, K., Gout, J.F., Kapusta, A., Betermier, M. and Meyer, E. (2011) Functional specialization of Piwi proteins in Paramecium tetraurelia from post-transcriptional gene silencing to genome remodelling. Nucleic Acids Res, 39, 4249–4264.

32. de Vanssay, A., Touzeau, A., Arnaiz, O., Frapporti, A., Phipps, J. and Duharcourt, S. (2020) The Paramecium histone chaperone Spt16-1 is required for Pgm endonuclease function in programmed genome rearrangements. PLoS Genet., 16, e1008949.

33. Arnaiz, O., Mathy, N., Baudry, C., Malinsky, S., Aury, J.-M., Denby Wilkes, C., Garnier, O., Labadie, K., Lauderdale, B.E., Le Mouël, A., et al. (2012) The Paramecium Germline Genome Provides a Niche for Intragenic Parasitic DNA: Evolutionary Dynamics of Internal Eliminated Sequences. PLoS Genet., 8, e1002984.

34. Arnaiz, O., Meyer, E. and Sperling, L. (2020) ParameciumDB 2019: integrating genomic data across the genus for functional and evolutionary biology. Nucleic Acids Res., 48, D599–D605.

35. Johnson, N.R., Yeoh, J.M., Coruh, C. and Axtell, M.J. (2016) Improved Placement of Multi-mapping Small RNAs. G3 *Bethesda* *Md*, **6**, 2103–2111.

36. Carradec, Q., Gotz, U., Arnaiz, O., Pouch, J., Simon, M., Meyer, E. and Marker, S. (2015) Primary and secondary siRNA synthesis triggered by RNAs from food bacteria in the ciliate Paramecium tetraurelia. Nucleic Acids Res., 43, 1818–1833.

37. Karunanithi, S., Oruganti, V., Marker, S., Rodriguez-Viana, A.M., Drews, F., Pirritano, M., Nordström, K., Simon, M. and Schulz, M.H. (2019) Exogenous RNAi mechanisms contribute to transcriptome adaptation by phased siRNA clusters in Paramecium. Nucleic Acids Res., 47, 8036–8049.

38. Solberg, T., Mason, V., Wang, C. and Nowacki, M. (2023) Developmental mRNA clearance by PIWI-bound endo-siRNAs in Paramecium. Cell Rep., 42, 112213.

39. Arnaiz, O., Cain, S., Cohen, J. and Sperling, L. (2007) ParameciumDB: a community resource that integrates the Paramecium tetraurelia genome sequence with genetic data. Nucleic Acids Res., 35, D439–444.

40. Zangarelli, C., Arnaiz, O., Bourge, M., Gorrichon, K., Jaszczyszyn, Y., Mathy, N., Escoriza, L., Betermier, M. and Regnier, V. (2022) Developmental timing of programmed DNA elimination in Paramecium tetraurelia recapitulates germline transposon evolutionary dynamics. Genome Res., 10.1101/gr.277027.122.

41. Nakanishi, K., Weinberg, D.E., Bartel, D.P. and Patel, D.J. (2012) Structure of yeast Argonaute with guide RNA. Nature, 486, 368–374.

42. Baumberger, N. and Baulcombe, D.C. (2005) Arabidopsis ARGONAUTE1 is an RNA Slicer that selectively recruits microRNAs and short interfering RNAs. Proc. Natl. Acad. Sci., 102, 11928–11933.

43. Liu, J., Carmell, M.A., Rivas, F.V., Marsden, C.G., Thomson, J.M., Song, J.-J., Hammond, S.M., Joshua-Tor, L. and Hannon, G.J. (2004) Argonaute2 Is the Catalytic Engine of Mammalian RNAi. Science, 305, 1437–1441.

44. Reuter, M., Berninger, P., Chuma, S., Shah, H., Hosokawa, M., Funaya, C., Antony, C., Sachidanandam, R. and Pillai, R.S. (2011) Miwi catalysis is required for piRNA amplification-independent LINE1 transposon silencing. Nature, 480, 264–267.

45. Rivas, F.V., Tolia, N.H., Song, J.-J., Aragon, J.P., Liu, J., Hannon, G.J. and Joshua-Tor, L. (2005) Purified Argonaute2 and an siRNA form recombinant human RISC. Nat. Struct. Mol. Biol., 12, 340–349.

46. Hoehener, C., Hug, I. and Nowacki, M. (2018) Dicer-like Enzymes with Sequence Cleavage Preferences. Cell, 173, 234–247.e7.

47. Matranga, C., Tomari, Y., Shin, C., Bartel, D.P. and Zamore, P.D. (2005) Passenger-strand cleavage facilitates assembly of siRNA into Ago2-containing RNAi enzyme complexes. Cell, 123, 607–620.

48. Miyoshi, K., Tsukumo, H., Nagami, T., Siomi, H. and Siomi, M.C. (2005) Slicer function of Drosophila Argonautes and its involvement in RISC formation. Genes Dev., 19, 2837–2848.

49. Rand, T.A., Petersen, S., Du, F. and Wang, X. (2005) Argonaute2 Cleaves the Anti-Guide Strand of siRNA during RISC Activation. Cell, 123, 621–629.

50. Noto, T., Kurth, H.M., Kataoka, K., Aronica, L., DeSouza, L.V., Siu, K.W.M., Pearlman, R.E., Gorovsky, M.A. and Mochizuki, K. (2010) The Tetrahymena Argonaute-Binding Protein Giw1p Directs a Mature Argonaute-siRNA Complex to the Nucleus. Cell, 140, 692–703.

51. Ignarski, M., Singh, A., Swart, E.C., Arambasic, M., Sandoval, P.Y. and Nowacki, M. (2014) Paramecium tetraurelia chromatin assembly factor-1-like protein PtCAF-1 is involved in RNA-mediated control of DNA elimination. Nucleic Acids Res., 42, 11952–11964.

52. Martinez, N.J. and Gregory, R.I. (2013) Argonaute2 expression is post-transcriptionally coupled to microRNA abundance. RNA, 19, 605–612.

53. Smibert, P., Yang, J.-S., Azzam, G., Liu, J.-L. and Lai, E.C. (2013) Homeostatic control of Argonaute stability by microRNA availability. Nat. Struct. Mol. Biol., 20, 789–795.

54. Bender, S., Tang, Y., Lindroth, A.M., Hovestadt, V., Jones, D.T.W., Kool, M., Zapatka, M., Northcott, P.A., Sturm, D., Wang, W., et al. (2013) Reduced H3K27me3 and DNA hypomethylation are major drivers of gene expression in K27M mutant pediatric high-grade gliomas. Cancer Cell, 24, 660– 672.

55. Lewis, P.W., Müller, M.M., Koletsky, M.S., Cordero, F., Lin, S., Banaszynski, L.A., Garcia, B.A., Muir, T.W., Becher, O.J. and Allis, C.D. (2013) Inhibition of PRC2 Activity by a Gain-of-Function H3 Mutation Found in Pediatric Glioblastoma. Science, 340, 857–861.

56. Lhuillier-Akakpo, M., Guérin, F., Frapporti, A. and Duharcourt, S. (2016) DNA deletion as a mechanism for developmentally programmed centromere loss. Nucleic Acids Res., 44, 1553–1565.

57. Gruchota, J., Denby Wilkes, C., Arnaiz, O., Sperling, L. and Nowak, J.K. (2017) A meiosis-specific Spt5 homolog involved in non-coding transcription. Nucleic Acids Res., 10.1093/nar/gkw1318.

58. Owsian, D., Gruchota, J., Arnaiz, O. and Nowak, J.K. (2022) The transient Spt4-Spt5 complex as an upstream regulator of non-coding RNAs during development. Nucleic Acids Res., 10.1093/nar/gkac106.

59. Jain, S.U., Rashoff, A.Q., Krabbenhoft, S.D., Hoelper, D., Do, T.J., Gibson, T.J., Lundgren, S.M., Bondra, E.R., Deshmukh, S., Harutyunyan, A.S., et al. (2020) H3 K27M and EZHIP Impede H3K27-Methylation Spreading by Inhibiting Allosterically Stimulated PRC2. Mol. Cell, 80, 726–735.e7.

60. Sarthy, J.F., Meers, M.P., Janssens, D.H., Henikoff, J.G., Feldman, H., Paddison, P.J., Lockwood, C.M., Vitanza, N.A., Olson, J.M., Ahmad, K., et al. (2020) Histone deposition pathways determine the chromatin landscapes of H3.1 and H3.3 K27M oncohistones. eLife, 9, e61090.

61. Brown, Z.Z., Müller, M.M., Jain, S.U., Allis, C.D., Lewis, P.W. and Muir, T.W. (2014) Strategy for “Detoxification” of a Cancer-Derived Histone Mutant Based on Mapping Its Interaction with the Methyltransferase PRC2. J. Am. Chem. Soc., 136, 13498–13501.

62. Stafford, J.M., Lee, C.-H., Voigt, P., Descostes, N., Saldaña-Meyer, R., Yu, J.-R., Leroy, G., Oksuz, O., Chapman, J.R., Suarez, F., et al. (2018) Multiple modes of PRC2 inhibition elicit global chromatin alterations in H3K27M pediatric glioma. Sci. Adv., 4, eaau5935.

63. Lee, J.M., Lee, J.S., Kim, H., Kim, K., Park, H., Kim, J.-Y., Lee, S.H., Kim, I.S., Kim, J., Lee, M., et al. (2012) EZH2 generates a methyl degron that is recognized by the DCAF1/DDB1/CUL4 E3 ubiquitin ligase complex. Mol. Cell, 48, 572–586.

64. He, A., Shen, X., Ma, Q., Cao, J., von Gise, A., Zhou, P., Wang, G., Marquez, V.E., Orkin, S.H. and Pu, W.T. (2012) PRC2 directly methylates GATA4 and represses its transcriptional activity. Genes Dev., 26, 37– 42.

65. Sanulli, S., Justin, N., Teissandier, A., Ancelin, K., Portoso, M., Caron, M., Michaud, A., Lombard, B., da Rocha, S.T., Offer, J., et al. (2015) Jarid2 Methylation via the PRC2 Complex Regulates H3K27me3 Deposition during Cell Differentiation. Mol. Cell, 57, 769–783.

66. Zhang, Q., Agius, S.C., Flanigan, S.F., Uckelmann, M., Levina, V., Owen, B.M. and Davidovich, C. (2021) PALI1 facilitates DNA and nucleosome binding by PRC2 and triggers an allosteric activation of catalysis. Nat. Commun., 12, 4592.

67. Ardehali, M.B., Anselmo, A., Cochrane, J.C., Kundu, S., Sadreyev, R.I. and Kingston, R.E. (2017) Polycomb Repressive Complex 2 Methylates Elongin A to Regulate Transcription. Mol. Cell, 68, 872–884.e6.

68. Margueron, R., Li, G., Sarma, K., Blais, A., Zavadil, J., Woodcock, C.L., Dynlacht, B.D. and Reinberg, D. (2008) Ezh1 and Ezh2 maintain repressive chromatin through different mechanisms. Mol. Cell, 32, 503–518.

69. Ji, L. and Chen, X. (2012) Regulation of small RNA stability: methylation and beyond. Cell Res., 22, 624–636.

70. Kurth, H.M. and Mochizuki, K. (2009) 2’-O-methylation stabilizes Piwi-associated small RNAs and ensures DNA elimination in Tetrahymena. Rna, 15, 675–85.

71. Han, J., LaVigne, C.A., Jones, B.T., Zhang, H., Gillett, F. and Mendell, J.T. (2020) A ubiquitin ligase mediates target-directed microRNA decay independently of tailing and trimming. Science, 370, eabc9546.

72. Shi, C.Y., Kingston, E.R., Kleaveland, B., Lin, D.H., Stubna, M.W. and Bartel, D.P. (2020) The ZSWIM8 ubiquitin ligase mediates target-directed microRNA degradation. Science, 370, eabc9359.

73. Ohtani, H., Iwasaki, Y.W., Shibuya, A., Siomi, H., Siomi, M.C. and Saito, K. (2013) DmGTSF1 is necessary for Piwi-piRISC-mediated transcriptional transposon silencing in the Drosophila ovary. Genes Dev., 27, 1656–1661.

74. Yoshimura, T., Watanabe, T., Kuramochi-Miyagawa, S., Takemoto, N., Shiromoto, Y., Kudo, A., Kanai-Azuma, M., Tashiro, F., Miyazaki, S., Katanaya, A., et al. (2018) Mouse GTSF1 is an essential factor for secondary piRNA biogenesis. EMBO Rep., 19, e42054.

75. Perez-Riverol, Y., Bai, J., Bandla, C., García-Seisdedos, D., Hewapathirana, S., Kamatchinathan, S., Kundu, D.J., Prakash, A., Frericks-Zipper, A., Eisenacher, M., et al. (2022) The PRIDE database resources in 2022: a hub for mass spectrometry-based proteomics evidences. Nucleic Acids Res., 50, D543– D552.

76. Berger, J.D. (1986) Autogamy in Paramecium. Cell cycle stage-specific commitment to meiosis. Exp Cell Res, 166, 475–85.

