## Supplementary Figures for "A histone methyltransferase-independent function of PRC2 controls small RNA dynamics during programmed DNA elimination in *Paramecium*"

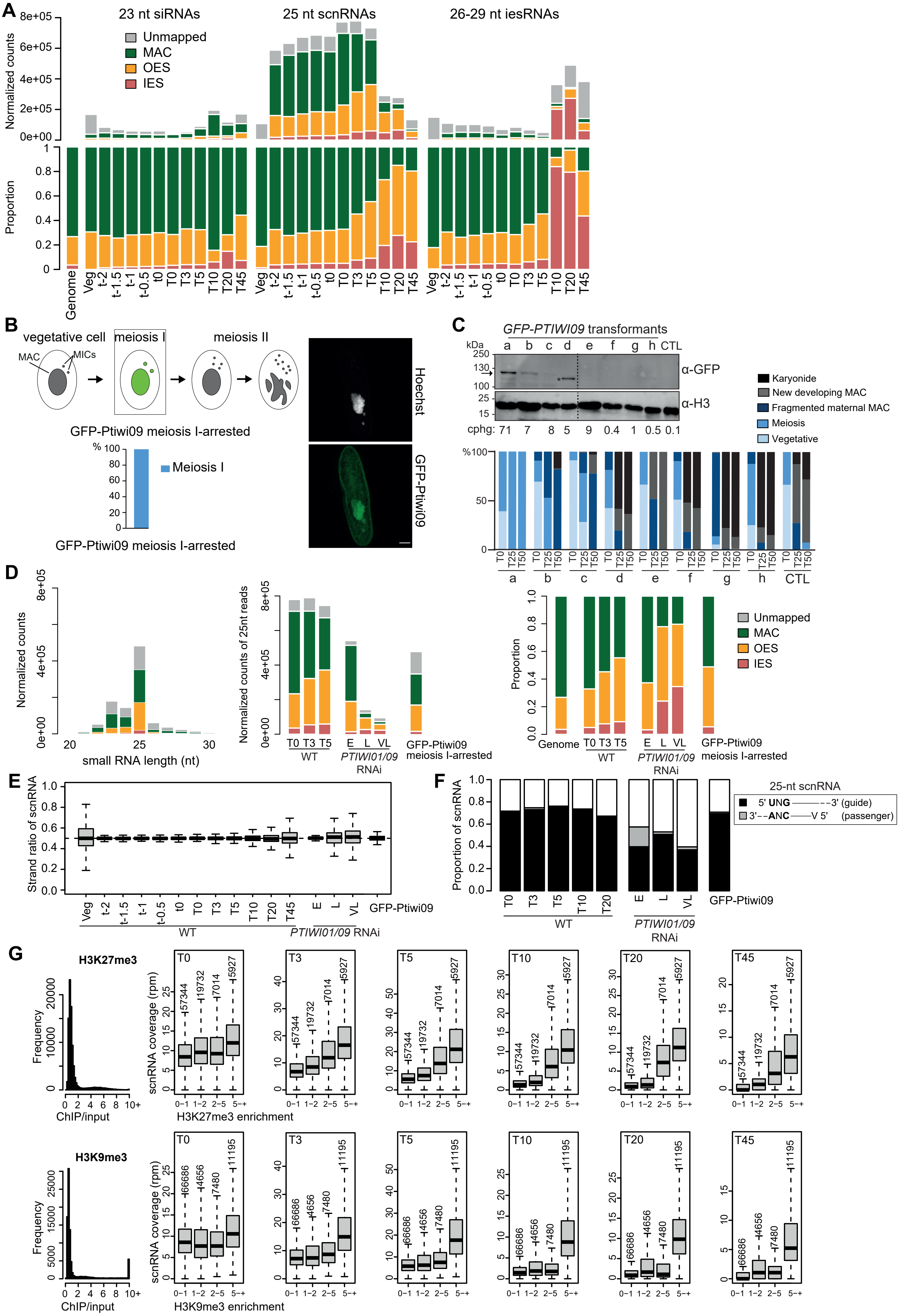

A
23 nt siRNAs
25 nt scnRNAs
26-29 nt iesRNAs
Unmapped
MAC
OES
IES
Normalized counts
Proportion
Genome
B
vegetative cell
meiosis I
meiosis II
MAC
MICs
GFP-Ptiwi09 meiosis I-arrested
Meiosis I
Hoechst
GFP-Ptiwi09
C
GFP-PTIWI09 transformants
α-GFP
α-H3
Karyonide
New developing MAC
Fragmented maternal MAC
Meiosis
Vegetative
D
Normalized counts
small RNA length (nt)
Normalized counts of 25nt reads
WT
PTIWI01/09 RNAi
GFP-Ptiwi09 meiosis I-arrested
E
Strand ratio of scnRNA
F
Proportion of scnRNA
25-nt scnRNA
5' UNG———3' (guide)
3'——ANC——V 5' (passenger)
G
H3K27me3
Frequency
ChIP/input
scnRNA coverage (rpm)
H3K27me3 enrichment
H3K9me3
H3K9me3 enrichment

**Figure S1. Related to Figure 2. sRNA dynamics during *Paramecium* sexual cycle.**

**A.** Analysis of sRNA populations (siRNAs; scnRNAs; iesRNAs) in vegetative cells and at different time points during autogamy. Bar plots show the normalized counts, using the total number of sRNA between 20 and 30-nt long, for each sample that map the MAC genome, IESs, or OES (top) and the proportion of reads for each category (middle). Genome: proportion of each category (MAC, OES, IES) in the MIC genome.

**B.** *GFP-PTIWI09* transformed cells arrested in meiosis I. Left: cytology of GFP-Ptiwi09 meiosis I-arrested. Right: GFP-Ptiwi09 localization in meiosis I-arrested cells. A confocal representative image is displayed. Overlay of Z-projections are presented. Scale bar is 10 µm.

**C.** Transformants with different copy numbers of the *GFP-PTIWI09* transgene (a to h). cphg: copy per haploid genome. Top: Western blot analysis from whole cells extracts at T20 with anti-GFP and -H3 antibodies. The arrow displays the Ptiwi09-GFP protein and the asterisk indicates an unspecific band. The dotted line indicates where the membrane has been cut (whole membranes in zenodo). Bottom: cytology at T0, T25 and T50 after the onset of autogamy. In the transformant with the highest copy number of the *GFP-PTIWI09* transgene (a), cells are arrested in meiosis I. At least 100 cells were scored for each time point by fluorescence microscopy with Hoechst staining.

**D.** Analysis of sRNA populations in GFP-Ptiwi09 meiosis-arrested cells. Left: Bar plots show the normalized reads at each sRNA size that are unmapped or match the MAC genome, IES and OES reference sequences. Middle: Normalized counts of 25-nt scnRNAs in wild-type cells at T0, T3 and T5, in *PTIWI01/09* RNAi conditions (Early (E), Late (L) and Very Late (VL) time points) (data from Furrer et al., 2017) and in GFP-Ptiwi09 meiosis I-arrested cells. Bar plots show the reads that are unmapped or match the MAC genome, IES and OES reference sequences. Right: Proportion of 25-nt scnRNAs that match the MAC genome, IES and OES reference sequences for the same samples. Genome: proportion of each category (MAC, OES, IES) in the MIC genome.

**E.** scnRNAs are produced from both strands of the MIC genome. For each time point, the boxplots display the distribution of the scnRNA strand ratio for the 189 longest scaffolds (MAC+IES genome assembly). A strand ratio of 0.5, indicated by the horizontal dotted line, means that 50% of the scnRNAs that mapped on the scaffold are transcribed in one orientation and 50% in the other.

**F.** Analysis of scnRNA composition signatures. The barplots show the proportion of 25-nt scnRNAs with the 5'-UNG signature characteristics of the guide strand (black), or those that do not start with a 5'-U (A, G or C: V) and end with a CNANN-3' signature, characteristics of the passenger strand (grey). scnRNA signatures for wild-type time course (T0 to T20), *PTIWI01/09* RNAi time points (from Furrer et al., 2017) and the GFP-Ptiwi09 meiosis I-arrested time point are presented.

**G.** Left: H3K27me3 (top) or H3K9me3 (bottom) enrichment (ChIP/input) upon *PGM* RNAi at T50 are computed over 1-kb bins. Right: ScnRNA coverage (RPM) at different time points of autogamy (T0 to T45) is shown as a function of H3K27me3 or H3K9me3 enrichment.

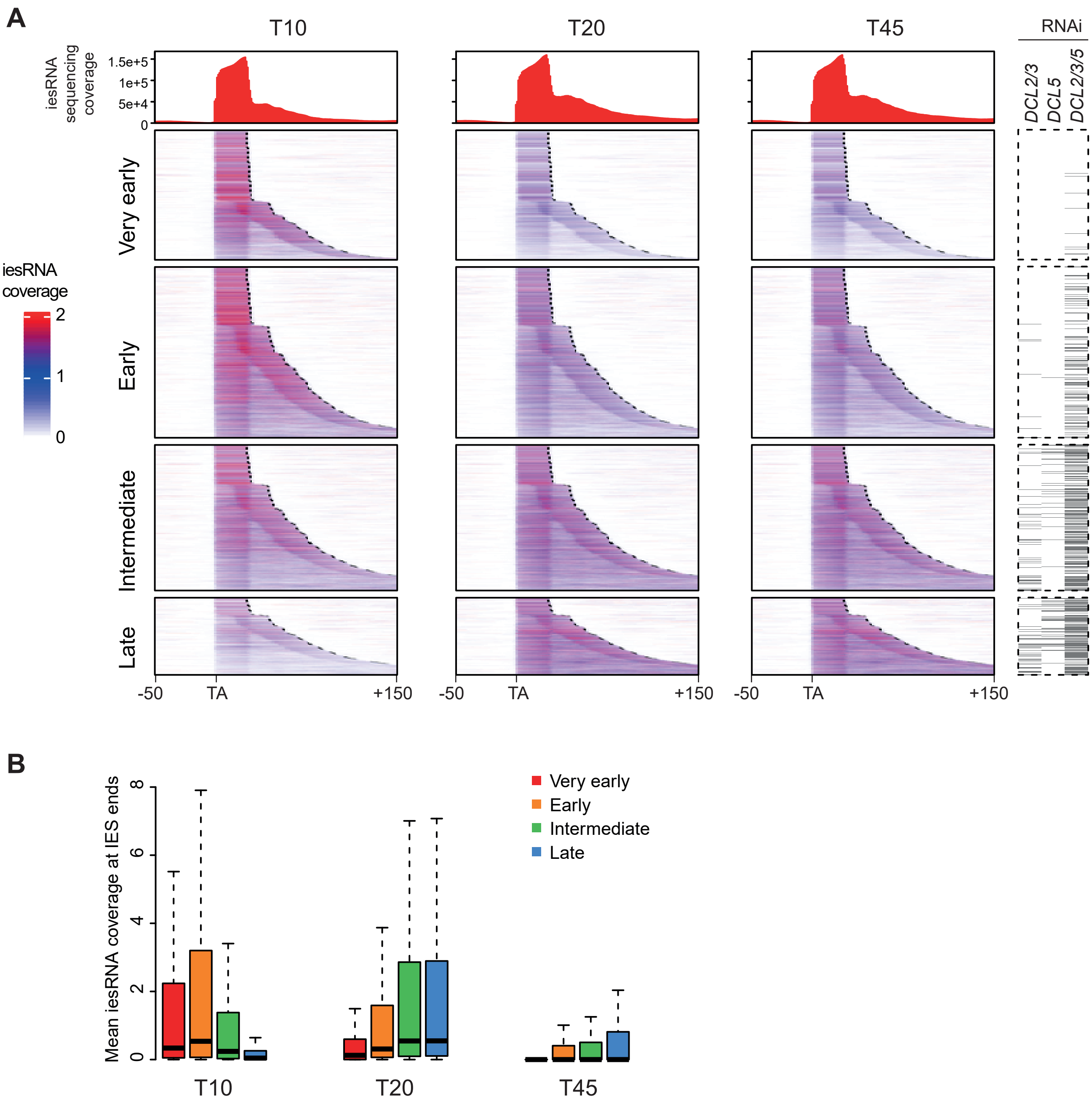


**Figure S2. Related to supplementary text. Localization of iesRNAs within IESs.**

**A.** At each time point (T10, T20, T45), the red histograms show the sequencing depth in iesRNAs around and inside all IESs. The TA dinucleotides at the left boudaries of IESs are aligned. The coverage is calculated 50 nt before and 150 nt after the TA (inside and outside the IES). The black dotted lines indicate the right boundaries of IESs. The heatmaps show, for the 4 excision timing IES groups (very early, early, intermediate and late), the position of iesRNAs in the genomic region of interest. The IESs, displayed in each line, are sorted according to their lengths in increasing order. The colorkey (white, blue, red) indicates the iesRNA coverage (log scale). On the right, the grey lines indicate the significantly *DCL2/3, DCL5* and *DCL2/3/5* -dependent IESs in the corresponding RNAi.

**B.** Boxplots show the mean iesRNA coverage distribution for the IESs belonging to each IES excision timing group.

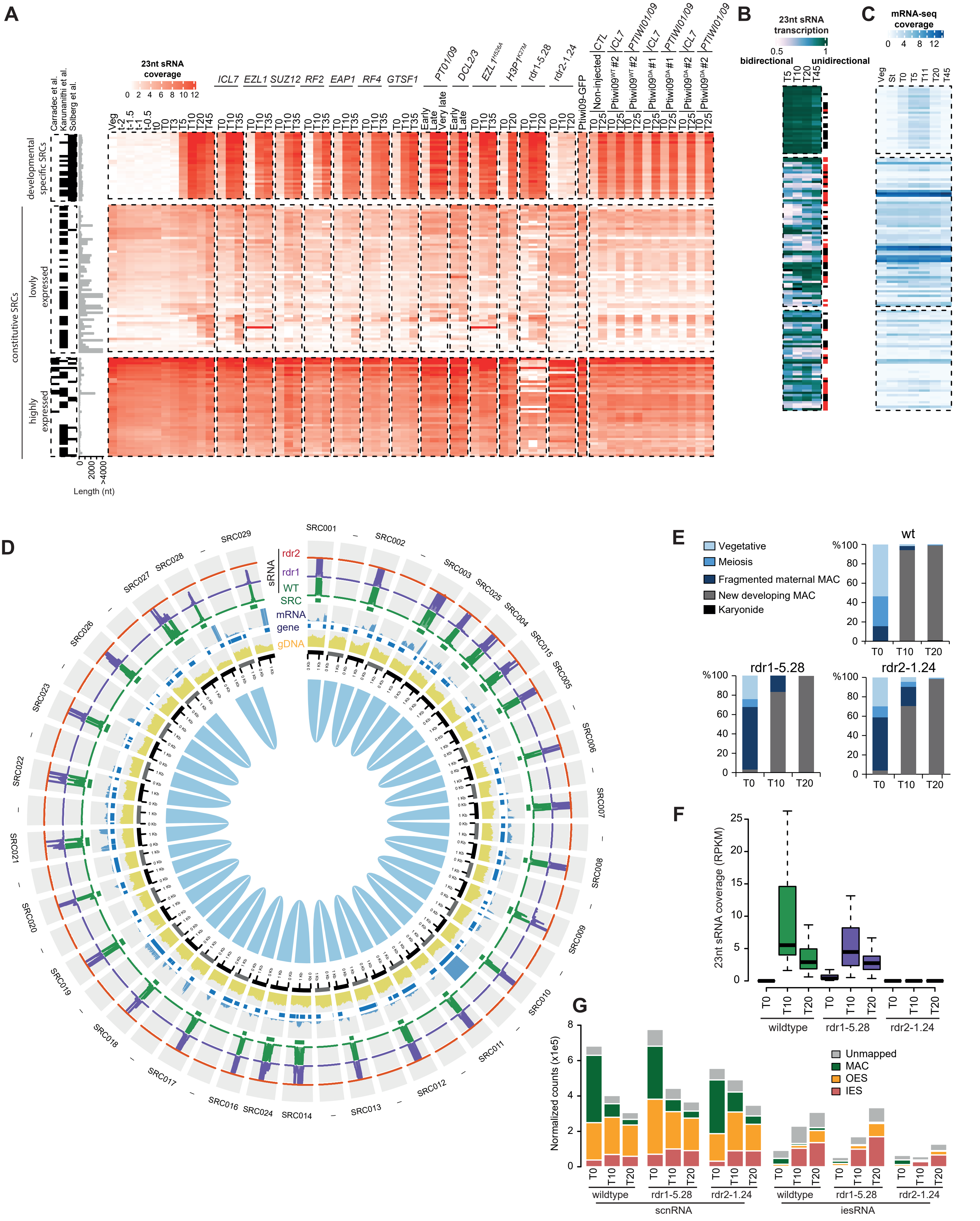
A
Carradec et al.
Karunanithi et al.
Solberg et al.
23nt sRNA coverage
0 2 4 6 8 10 12
ICL7
EZL1
SUZ12
RF2
EAP1
RF4
GTSF1
PT01/09
DCL2/3
EZL1 H526A
H3P1 K27M
rdr1-5.28
rdr2-1.24
Ptiwi09-GFP
CTL
Non-injected
ICL7
PTIWI01/09
Veg
t-2
t-1.5
t-1
t-0.5
t0
T0
T3
T5
T10
T20
T35
T45
Early
Late
Very late
Ptiwi09WT #2
Ptiwi09DA #1
Ptiwi09DA #2
developmental specific SRCs
constitutive SRCs
lowly expressed
highly expressed
0
2000
>4000
Length (nt)
B
23nt sRNA transcription
0.5
1
bidirectional
unidirectional
T5
T10
T20
T45
C
mRNA-seq coverage
0 4 8 12
Veg
St
T0
T5
T11
T20
T45
D
sRNA
rdr2
rdr1
WT
SRC
mRNA
gene
gDNA
SRC001
SRC002
SRC003
SRC025
SRC004
SRC015
SRC005
SRC006
SRC007
SRC008
SRC009
SRC010
SRC011
SRC012
SRC013
SRC014
SRC024
SRC016
SRC017
SRC018
SRC019
SRC020
SRC021
SRC022
SRC023
SRC026
SRC027
SRC028
SRC029
E
Vegetative
Meiosis
Fragmented maternal MAC
New developing MAC
Karyonide
wt
rdr1-5.28
rdr2-1.24
%100
80
60
40
20
0
T0
T10
T20
F
23nt sRNA coverage (RPKM)
25
20
15
10
5
0
wildtype
rdr1-5.28
rdr2-1.24
G
Normalized counts (x1e5)
8
6
4
2
0
Unmapped
MAC
OES
IES
scnRNA
iesRNA

**Figure S3. Related to supplementary text. Developmental-specific 23 nt short-read clusters (SRCs).**

**A.** The heatmaps display the normalized 23-nt sRNA coverage (log2 RPKM calculated using the number of sRNA between 20 and 30-nt long) for the 136 SRCs during autogamy time-course experiments. SRCs were split using a K-means clustering in 3 groups: developmental-specific SRCs (N=29), lowly constitutively expressed SRCs (N=64) and highly constitutively expressed SRCs (N=43) (see Materials Methods). On the left, the black lines indicate the SRCs identified in previous studies (Carradec et al., 2015; Karunanithi et al., 2019; Solberg et al., 2024). Length distribution of the 136 SRCs and overlap with previous studies are indicated. In addition to the WT autogamy time course data, the heatmap shows the normalized 23-nt coverage under different conditions and at different time-points. Note that depletion of PRC2-Ezl1 core components (Ezl1, Suz12) and associated cofactors (Rf2, Eap1, Rf4), of Gtsf1, or expression of the Ptiwi09 slicer mutant had no apparent impact on the accumulation of 23-nt developmental-specific SRCs.

**B.** Heatmap showing the 23-nt sRNA transcription orientation for the 136 SRCs. White indicates a 50% bidirectional transcription ratio and green a unidirectional transcription ratio. The SRCs which overlap a putative coding gene are indicated in black when they are in antisense orientation, and in red when in the same orientation as its overlapping gene.

**C.** Heatmap showing the normalized mRNA-seq coverage (log2 RPKM) (data from PRJEB19343 in Arnaiz et al., 2017). “St” corresponds to “meiosis” stage.

**D.** Circos representation of the regions around of the 29 developmental-specific SRCs (SRC001 to SRC029) and their paralogous genomic regions issued from the most recent WGD (blue arcs). SRC names are displayed when paralogous regions correspond to SRCs. The SRCs (green circles) are localized on genomic regions defined by 500 nt upstream and downstream SRC boundaries (black and grey circles). Blue circles represent the part of the regions covered by genes. Histograms represent the sequencing coverage: genomic DNA (yellow), mRNA coverage (at T10 and T20, blue), and 23-nt sRNA coverage (at T0 and T20) in wild-type (wt) condition (green), rdr1 mutant (purple) and rdr2 mutant (orange).

**E.** Progression of autogamy is followed by cytology with Hoechst staining in each condition (wild-type, rdr1-5.28 mutant, rdr2-1.24 mutant). At least 100 cells were scored for each time point by fluorescence microscopy.

**F.** Boxplot showing global the normalized 23-nt sRNA coverage (RPKM calculated using the number of sRNA between 20 and 30-nt long) distribution for developmental-specific SRCs at 3 time points (T0, T10 and T20) in 3 conditions: wild-type, rdr1-5.28 mutant and rdr2–1.24 mutant.

**G.** Analysis of sRNA populations (scnRNAs and iesRNAs) at different time points during autogamy in wildtype condition, rdr1-5.28 mutant and rdr2–1.24 mutant. Bar plots show the normalized counts, using the total number of sRNA between 20 and 30-nt long, for each sample that map the MAC genome, IES, OES or unmapped on the MIC genome.

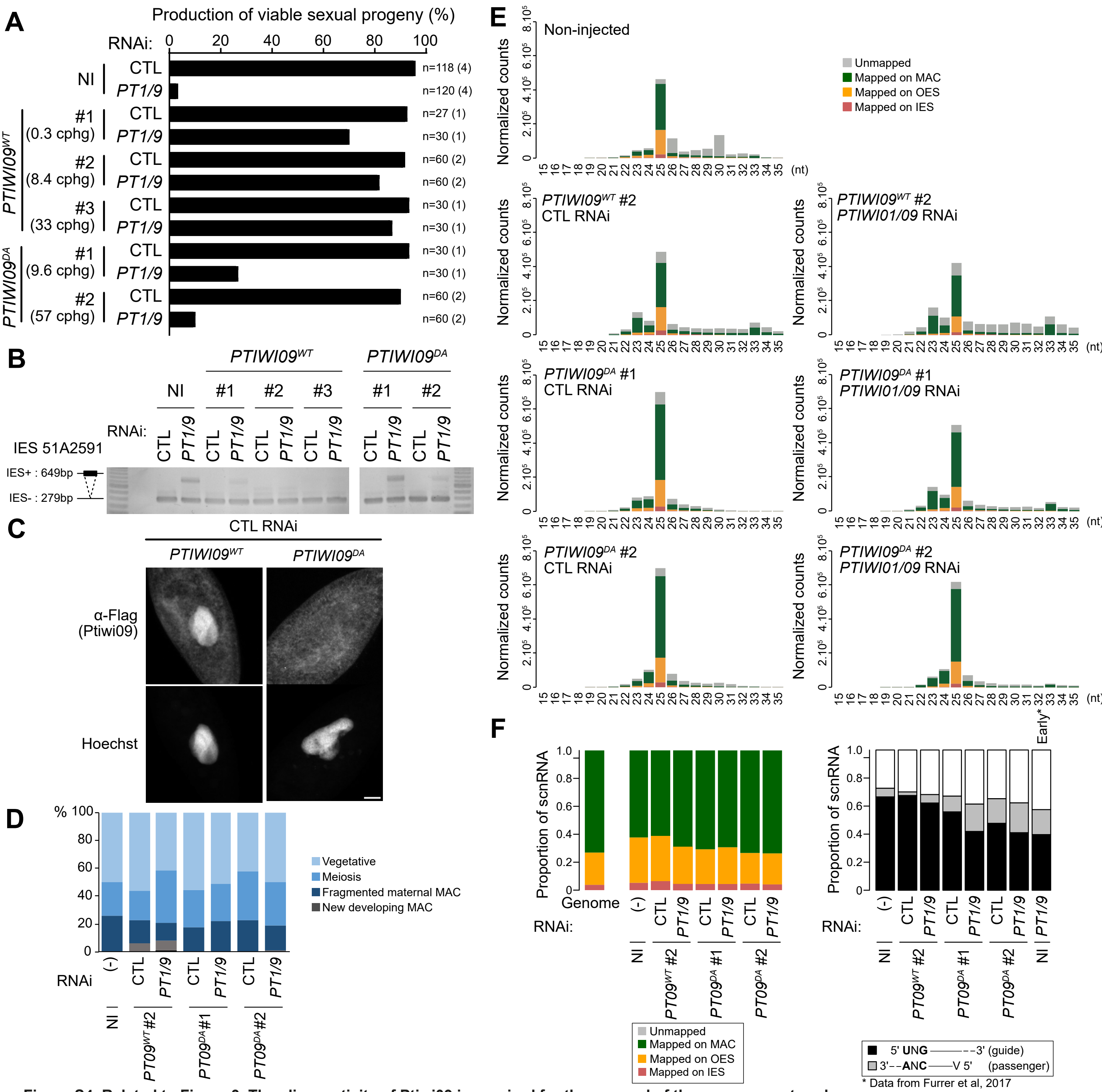


**Figure S4. Related to Figure 3. The slicer activity of Ptiwi09 is required for the removal of the passenger strand.**

**A.** Production of viable sexual progeny of non-injected (NI) cells and cells transformed with *3XFLAG-HA-PTIWI09* (*PTIWI09*$^{WT}$) or *3XFLAG-HA-PTIWI09*$^{DA}$ (*PTIWI09*$^{DA}$), following *PTIWI01/09* (*PT1/9*) or *ICL7* (CTL) RNAi. For each transformant, the number of transgene copy per haploid genome (cphg) is indicated in parentheses. The total number of cells analyzed for each RNAi and the number of independent experiments (in parentheses) are indicated at the end of each bar. Related to Figure 3A.

**B.** PCR analysis of IES retention with primers located on either side of IES 51A2591 in non-injected (NI) cells and cells transformed with 3*XFLAG-HA-PTIWI09* (*PTIWI09*$^{WT}$) or *3XFLAG-HA-PTIWI09*$^{DA}$ (*PTIWI09*$^{DA}$), in *ICL7* (CTL) and *PTIWI01/09* (*PT1/9*) RNAi conditions. Total DNA samples were prepared from post-autogamous cells. Because the maternal MAC is still present at this stage, the excised version is amplified in all cases; the IES-retaining fragment can only be detected if present in the developing MACs. Related to Figure 3A.

**C.** FLAG immunostaining at T0 of cells transformed with *3XFLAG-HA-PTIWI09* (*PTIWI09*$^{WT}$ #2) or *3XFLAG-HA-PTIWI09*$^{DA}$ (*PTIWI09*$^{DA}$ #2), following *ICL7* (CTL) RNAi-mediated silencing. Representative confocal images are displayed. Overlay of Z-projections are presented. Scale bar is 10 µm. Related to Figure 3C.

**D.** Description of the cytological stages of the samples used for sRNAseq. Progression of autogamy was followed by cytology with Hoechst staining in non-injected (NI) cells without RNAi (-), and cells transformed with *3XFLAG-HA-PTIWI09* (*PTIWI09*$^{WT}$) or *3XFLAG-HA-PTIWI09*$^{DA}$ (*PTIWI09*$^{DA}$), in *ICL7* (CTL) and *PTIWI01/09* (*PT1/9*) RNAi conditions. At least 100 cells were scored for each condition by fluorescence microscopy. Related to Figure 3D.

**E.** Analysis of sRNA populations at T0 in non-injected (NI) cells without RNAi, and cells transformed with *3XFLAG-HA-PTIWI09* ( *PTIWI09*$^{WT}$) or *3XFLAG-HA-PTIWI09*$^{DA}$ ( *PTIWI09*$^{DA}$), in *ICL7* (CTL) and *PTIWI01/09* RNAi conditions. Bar plots show the normalized reads at each sRNA size that are unmapped, or match the MAC genome, IES and OES reference sequences. Related to Figure 3D.

**F.** Analysis of 25-nt scnRNAs at T0 in non-injected (NI) cells and cells transformed with *3XFLAG-HA-PTIWI09* ( *PTIWI09*$^{WT}$) or *3XFLAG-HA-PTIWI09*$^{DA}$ (*PTIWI09*$^{DA}$), without RNAi (-), in *ICL7* (CTL) and *PTIWI01/09* (*PT1/9*) RNAi conditions. Left: Proportion of 25-nt scnRNAs that match the MAC genome, IESs, and OES reference sequences. Genome: proportion of each category (MAC, OES, IES) in the MIC genome. Right: Bar plots show the proportion of 25-nt scnRNAs with the 5'-UNG signature, characteristics of the guide strand (black), or those that do not start with a 5'-U (A, G or C: V) and end with a CNANN-3' signature, characteristics of the passenger strand (grey). Data for NI, *PTIWI09*$^{WT}$ #2 and *PTIWI09*$^{DA}$ #2 are shown in Figure 3D.

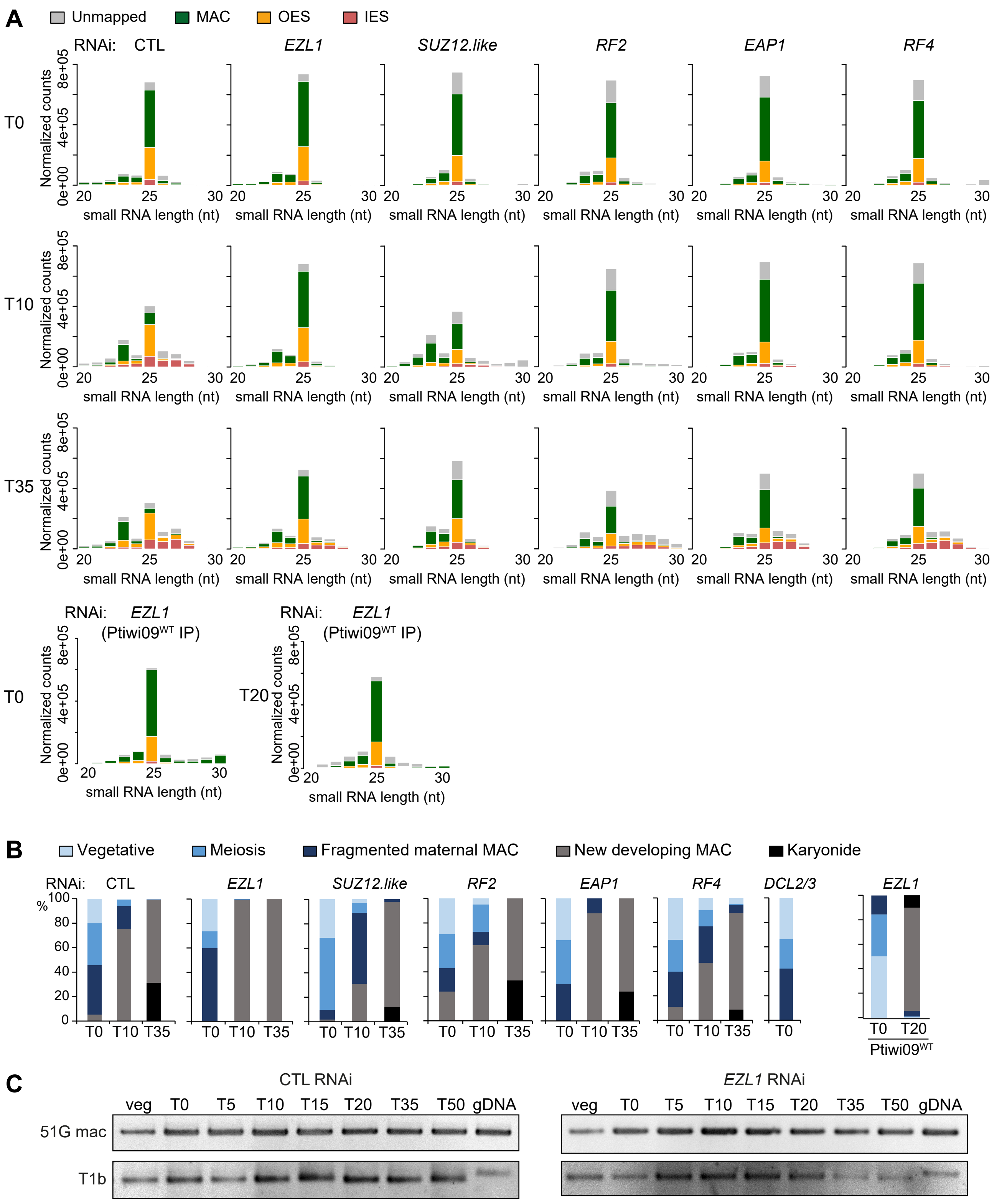


**Figure S5. Related to Figure 4. PRC2-Ezl1 core complex and its associated cofactors are required for scnRNA selection.**

**A.** Analysis of small RNA populations at T0, T10 and T35 in ICL7 (CTL), *EZL1, SUZ12.like*, *RF2, EAP1* and *RF4* RNAi conditions related to Figure 4A-B), at T0 and T20 in *EZL1* RNAi after Ptiwi09 immunoprecipitation. Bar plots show the normalized reads at each sRNA size that are unmapped or match the MAC genome, IESs and OES reference sequences.

**B.** Description of the cytological stages of the samples used for sRNAseq. The *DCL2/3* RNAi sample is described in Sandoval et al. 2014. Progression of autogamy was followed by cytology with Hoechst staining in the time course experiments upon RNAi-mediated KD. At least 100 cells were scored for each time point by fluorescence microscopy.

**C.** Detection of 51G mac ncRNA transcripts by RT-PCR in control (CTL) and *EZL1* RNAi conditions. Total RNAs, extracted at each time point, were reverse-transcribed and cDNAs were amplified by PCR with gene-specific primers and, as a loading control, with primers on either side of an intron for the constitutively-expressed gene encoding a trichocyst matrix protein (T1b). Genomic DNA (gDNA) was used as a positive PCR control. Cell cultures do not express the 51G surface antigen gene. Transcripts of the 51G gene that can be detected by RT–PCR both during vegetative growth and during sexual events are non-coding RNAs expressed from the MAC, as previously reported (Lepere et al., 2008; Maliszewska-Olejniczak et al., 2015).

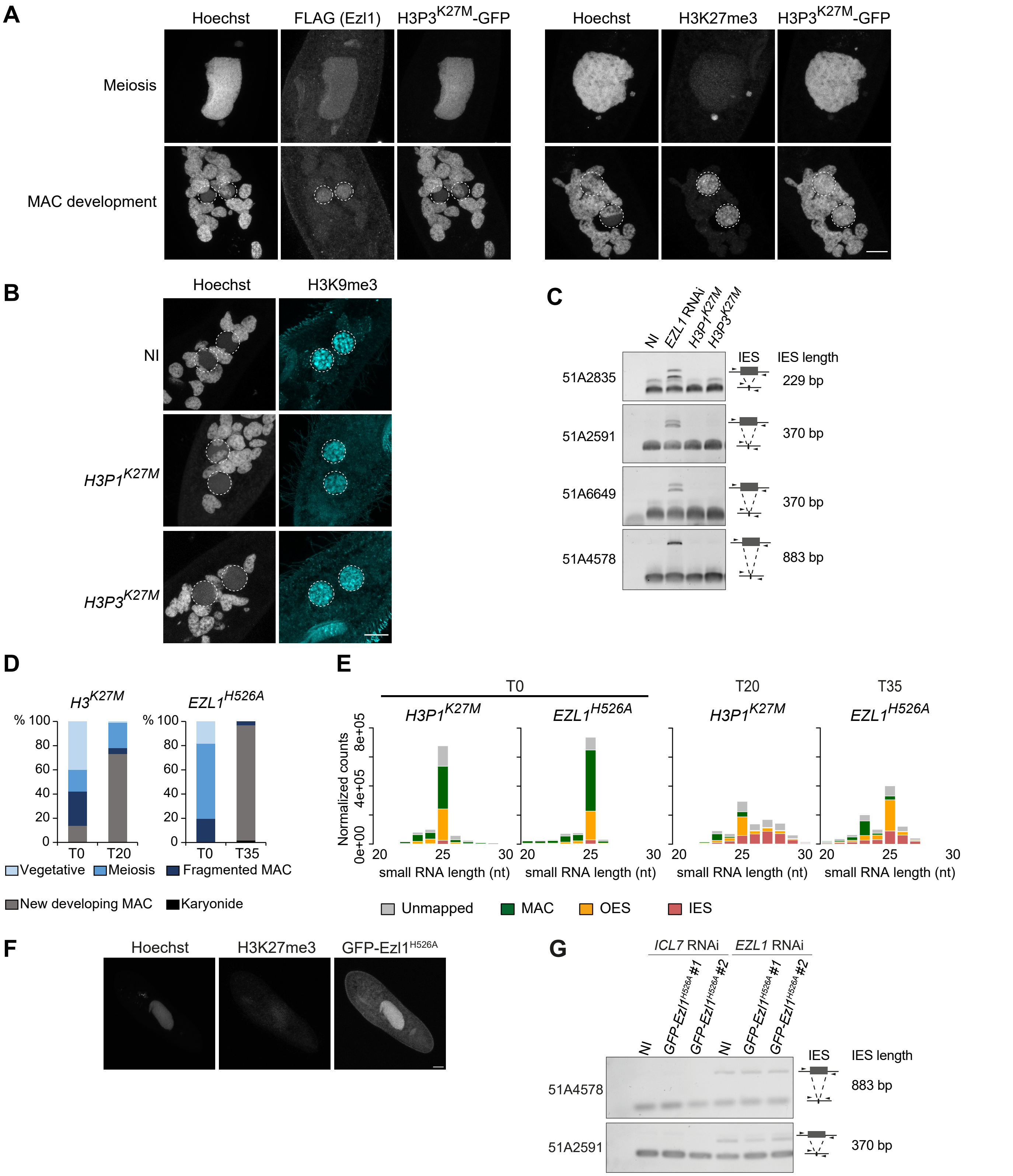


**Figure S6. Related to Figure 5. Ezl1 histone methyltransferase activity is not required for scnRNA selection.**

**A.** GFP fluorescence and FLAG or H3K27me3 immunostaining of cells transformed with H3P3$^{K27M}$-*GFP* and *3XFLAG-HA-EZL1* at MIC meiosis or during new MAC development. Z-projections of magnified views are presented. Dashed white circles indicate the new developing MACs. Scale bar is 10 µm.

**B.** H3K9me3 immunostaining of non-injected cells (NI), and of cells transformed with *H3P1*$^{K27M}$ or *H3P3*$^{K27M}$ during MAC development. Dashed white circles indicate the new developing MACs. Scale bar is 10 µm.

**C.** PCR analysis of IES retention with primers located on either side of the IES in cells transformed with *H3P1* or *H3P3* transgenes bearing K27M mutations (*H3P1*$^{K27M}$ or *H3P3*$^{K27M}$). Total DNA samples were prepared from starved post-autogamous cells. Because the maternal MAC is still present at this stage, the excised version is amplified in all cases; the IES-retaining fragment can only be detected if present in the developing MACs.

**D.** Description of the cytological stages of the samples used for sRNAseq.

**E.** Analysis of small RNA populations in cells transformed with *H3P1*$^{K27M}$ or with *GFP-EZL1*$^{H526A}$ upon endogenous *EZL1* RNAi. Bar plots show the normalized reads at each sRNA size that are unmapped or match the MAC genome, IESs and OES reference sequences.

**F.** H3K27me3 immunostaining and *GFP-EZL1*$^{H526A}$ localization in the maternal MAC during meiosis in cells transformed with *GFP-EZL1*$^{H526A}$ upon endogenous *EZL1* RNAi. Representative confocal images are displayed. Overlay of Z-projections of magnified views of Hoechst staining, H3K27me3-specific antibodies and GFP signal are presented. Scale bar is 10 µm.

**G.** PCR analysis of IES retention with primers located on either side of the IES in cells transformed with *GFP-EZL1*$^{H526A}$ transgenes for transformants #1 et #2. Total DNA samples were prepared from starved post-autogamous cells. Because the maternal MAC is still present at this stage, the excised version is amplified in all cases; the IES-retaining fragment can only be detected if present in the developing MACs. Expression of the GFP-Ezl1 fusion protein impairs IES elimination.

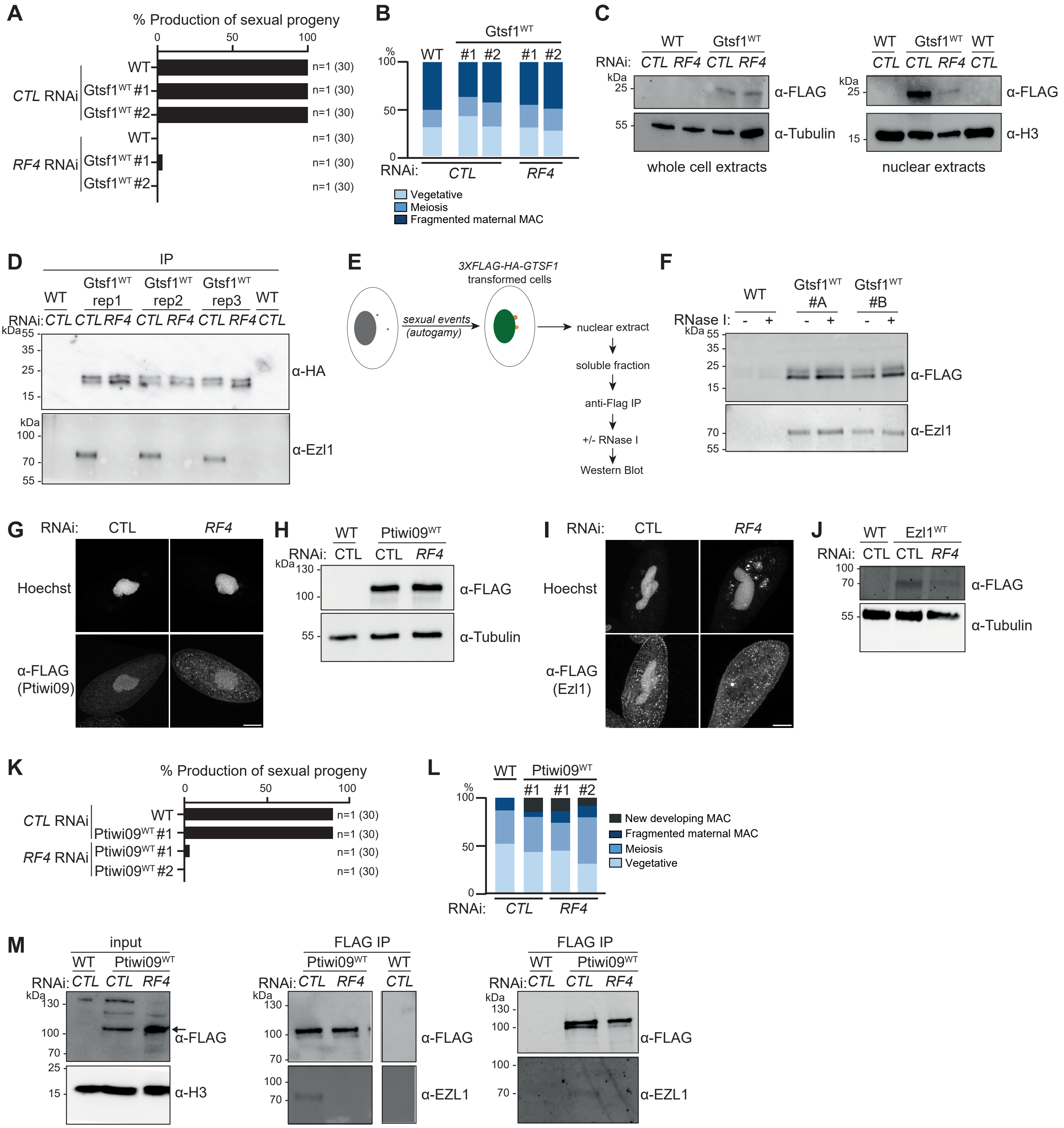
A
% Production of sexual progeny
0
50
100
CTL RNAi
RF4 RNAi
WT
Gtsf1WT #1
Gtsf1WT #2
n=1 (30)
B
Gtsf1WT
%
100
50
0
WT
#1
#2
RNAi:
CTL
RF4
Vegetative
Meiosis
Fragmented maternal MAC
C
WT
Gtsf1WT
RNAi:
CTL
RF4
kDa
25
55
15
α-FLAG
α-Tubulin
α-H3
whole cell extracts
nuclear extracts
D
IP
rep1
rep2
rep3
35
70
100
α-HA
α-Ezl1
E
3XFLAG-HA-GTSF1
transformed cells
sexual events
(autogamy)
nuclear extract
soluble fraction
anti-Flag IP
+/- RNase I
Western Blot
F
RNase I:
#A
#B
G
Hoechst
α-FLAG
(Ptiwi09)
H
Ptiwi09WT
130
I
(Ezl1)
J
Ezl1WT
K
Ptiwi09WT #1
Ptiwi09WT #2
L
New developing MAC
M
input
FLAG IP
α-EZL1

**Figure S7. Related to Figure 6. The PRC2 cofactor Rf4 mediates the interaction between Ptiwi09 and Gtsf1.**

**A.** Production of viable sexual progeny of wild-type cells (WT) and of two transformants (#1 and #2) expressing a 3xFLAG-HA-Gtsf1 (Gtsf1$^{WT}$) fusion protein following *ICL7* (CTL) or *RF4* RNAi (related to Figure 6A-B). The total number of cells analyzed for each RNAi and the number of independent experiments (in parentheses) are indicated. As expected, no viable sexual progeny is produced in *RF4* RNAi conditions.

**B.** Description of the cytological of the Gtsf1 IP experiments upon *ICL7* (CTL) or *RF4* RNAi (related to Figure 6A-B).

**C.** Western blot analyses of whole cell and nuclear extracts at T0 from non-injected cells (WT) and cells expressing a 3XFLAG-HA-Gtsf1 functional protein (Gtsf1) upon *ICL7* (CTL) or *RF4* RNAi. Anti-FLAG, anti-tubulin and anti-H3 antibodies are used for detection.

**D.** Gtsf1 immunoprecipitation replicates of the experiment shown in Figure 6B. FLAG immunoprecipitation was performed from nuclear extracts of non-injected cells (WT) and cells expressing a 3xFLAG-HA-Gtsf1 functional protein (Gtsf1$^{WT}$) upon *ICL7* (CTL) or *RF4* RNAi in 3 replicates in each RNAi (rep1; rep2; rep3). Western blot analysis of Gtsf1 and Ezl1 after affinity purification (IP). Anti-HA antibodies are used to detect Gtsf1, and anti-Ezl1 to detect Ezl1.

**E.** RNase1 treatment of Gtsf1 immunoprecipitation samples obtained from nuclear extracts of cells expressing a 3xFLAG-HA-Gtsf1 functional protein. After FLAG affinity purification, samples were treated with RNaseI or not, before western blot analysis.

**F.** Western blot analysis of the FLAG immunoprecipitation from nuclear extracts of non-injected cells (WT) or cells expressing a 3xFLAG-HA-Gtsf1 functional protein (Gtsf1$^{WT}$) for two replicates (#A and #B), with no treatment or RNase1 treatment.

**G.** Anti-FLAG immunostaining of cells transformed with a *3XFLAG-HA-PTIWI09* transgene upon *ICL7* (CTL) or *RF4* RNAi at T0. Representative confocal images are displayed. Overlay of Z-projections are presented. Scale bar is 20 µm.

**H.** Western blot analysis of whole cell extracts at T0 from non-injected cells (WT) and cells expressing a 3xFLAG-HA-Ptiwi09 fusion protein upon *ICL7* (CTL) or *RF4* RNAi. Anti-FLAG and anti-H3 antibodies are used for detection.

**I.** Anti-FLAG immunostaining of cells transformed with a *3XFLAG-HA-EZL1* transgene upon *ICL7* (CTL) or *RF4* RNAi at T0. Representative confocal images are displayed. Overlay of Z-projections are presented. Scale bar is 10 µm.

**J.** Western blot analysis of whole cell extracts at T0 from non-injected cells (WT) and cells expressing a 3xFLAG-HA-Ezl1 fusion protein upon *ICL7* (CTL) or *RF4* RNAi. Anti-FLAG and anti-tublin antibodies are used for detection.

**K.** Production of viable sexual progeny of wild-type cells (WT) and of two transformants (#1 and #2) expressing a 3xFLAG-HA-Ptiwi09 (Ptiwi09$^{WT}$) fusion protein following no RNAi (CTL), or *RF4* RNAi (related to Figure 6C-D). The total number of cells analyzed for each RNAi and the number of independent experiments (in parentheses) are indicated. As expected, no viable sexual progeny is produced in *GTSF1* or *RF4* RNAi conditions.

**L.** Description of the cytological of the Ptiwi09 IP experiments upon no RNAi (CTL) or *RF4* RNAi (related to Figure 6C-D).

**M.** Ptiwi09 immunoprecipitation replicates of the experiment shown in Figure 6D. FLAG immunoprecipitation was performed from nuclear extracts of non-injected cells (WT) and cells expressing a 3xFLAG-HA-Ptiwi09 functional protein (Ptiwi09$^{WT}$) upon no RNAi (CTL) or *RF4* RNAi. Western blot analysis of Ptiwi09 before (input) and after affinity purification (FLAG IP) for 2 additional replicates besides the one displayed in Figure 6D. Anti-FLAG antibodies are used to detect Ptiwi09, anti-Ezl1 to detect Ezl1, and H3 for normalization. The uncropped membranes can be found in zenodo.

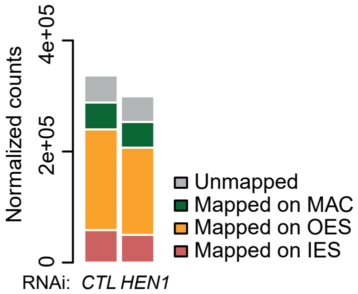


**Figure S8. scnRNA population upon *HEN1* RNAi**

Analysis of scnRNA populations at a late stage during autogamy in cells depleted for *ICL7* (CTL) of *HEN1* (Solberg et al., 2023). Bar plots show the normalized counts for each sample that map the MAC genome, IES, OES or unmapped on the MIC genome.

**Supplementary Text**

We took advantage of the sRNA dataset we had generated to examine two other small RNA populations during autogamy: the 26-29-nt long iesRNAs and the 23-nt long siRNAs.

We analyzed the sequencing data from all time points and mapped the iesRNAs on the MIC reference genome (see Methods). iesRNAs accumulated at T10. They mostly mapped IESs (Figure S1A), and the highest iesRNA coverage is found at the extremities of IESs (Figure S2A), as previously reported (Sandoval et al., 2014). We analyzed iesRNA coverage as a function of IES elimination timing (Figure S2A-B) (Zangarelli et al., 2022). We found that iesRNA accumulation correlates with IES elimination timing and all IESs produce iesRNAs, whether they are excised early or late (Figure S2A-B). The biogenesis of iesRNAs has been shown to require the Dcl5 protein, while scnRNA production depends on the Dcl2 and Dcl3 proteins (Lepere et al., 2009; Sandoval et al., 2014). Very early excised IESs are not sensitive to Dcl5 depletion, even though they are covered by iesRNAs (Figure S2A). Therefore, we conclude that all IESs can produce iesRNAs, regardless of whether their excision depends on Dcl2/3, Dcl5, or both (Figure S2A). Similarly, scnRNAs are produced for all IESs (Figure 2C), although only a subset of IESs require Dcl2/3 for their excision (Lhuillier-Akakpo et al., 2014; Sandoval et al., 2014).

Although 23-nt siRNAs are present at low levels during most stages of the life cycle, their levels increase slightly during autogamy (Figure S1A). To investigate whether this increase is due to the production of siRNAs from specific genomic loci, we analyzed the sequencing data from all time points and mapped the 23-nt siRNAs on the MAC genome (see Methods). Based on sequencing coverage (see Methods), we identified 136 short-read clusters (SRCs) (Tables S2-S3) producing siRNAs from specific genomic loci. As expected, 85 overlapped with previously described SRCs expressed constitutively during vegetative growth (Carradec et al., 2015; Karunanithi et al., 2019). 29 overlapped with recently reported development-specific SRCs, which produce siRNAs that are Dicer 1-dependent and are bound to the Ptiwi08 protein (Solberg et al., 2023) (Figure S3). These 29 developmental-specific SRCs, whose mean length is 451 nt, accumulated at T5 and were still detected at T45 (Figure S3A). During this time frame, the developmental-specific SRCs accounted for the majority of 23-nt siRNAs (79% at T10 mapped on the 29 developmental-specific SRCs). While scnRNAs are produced from both strands, siRNAs corresponding to these 29 SRCs were exclusively transcribed from one strand (Figure S3B). Because the 16 SRCs overlap with predicted genes, we examined the levels of messenger RNAs (Figure S3C). mRNA expression was detected at T0, earlier than that of SRCs, and with an apparent peak at T5 and T11 and low expression levels at T45. Overall, the mRNA expression profiles of SRCs appear not to correlate with the timing of siRNA expression emanating from SRCs. The fact that the predicted overlapping genes are smaller (median gene length 256 nt) than the average gene length (median gene length 1046 nt for all genes) raises the question of whether these RNAseq covered genes are lowly expressed coding genes or possibly pseudogenes. Moreover, for the 16 developmental-specific SRCs overlapping putative protein coding genes, we noted that sRNAs accumulated antisense to gene transcription for most of them (15/16) (Figure S3B), consistent with the idea that developmental-specific SRCs are not degradation products of gene messenger RNAs.
The *P. tetraurelia* genome has undergone several whole genome duplications (WGDs) in the course of evolution, the last one dated before the radiation of the aurelia species complex

(Aury et al., 2006; Sellis et al., 2021). We retrieved the paralogous region stemming from the last WGD for each developmental-specific SRC (Figure S3D). We identified paralogous SRCs for some of them (6/29) which indicate that those correspond to conserved SRCs. In contrast, in the other cases, the paralogous regions were either lost or no SRC could be detected (23/29), indicating that these SRCs may have been either acquired more recently, after the last WGD, or lost on the paralogous region.
To identify factors involved in the biogenesis of developmental-specific SRCs, we focused on two RNA-dependent RNA polymerases (Rdr1 and Rdr2) known to control the accumulation of endogenous siRNAs. Rdr2 was previously shown to be involved in the production of endogenous siRNAs (Marker et al., 2014) and a strong reduction in the accumulation of siRNAs from a few constitutive SRCs was reported with the rdr2-1.24 allele (Carradec et al., 2015) (Figure S3E). A null mutant of *RDR1* (rdr1-5.28 allele) also impaired accumulation of endogenous siRNAs (Carradec et al., 2015; Karunanithi et al., 2019) (Figure S3E). We sequenced the small RNA populations in these two rdr1 and rdr2 mutants at three time points (T0; T10 and T20) during an autogamy time course experiment (Figure S1A). While the rdr1 mutant had no impact on the accumulation of developmental-specific SRCs, accumulation of siRNAs from the developmental-specific SRCs was completely abolished in the rdr2 mutant (Figures S3A-S3D-S3E). We note that rdr1 and rdr2 mutants do not affect scnRNA and iesRNA biogenesis, nor do they have an impact on scnRNA selection (Figure S3G). In addition, we noted that depletion of PRC2-Ezl1 core components and associated cofactors had no apparent impact on the accumulation of 23-nt developmental-specific SRCs (Figure S3B).

In conclusion, we showed that accumulation of siRNAs from the developmental-specific SRCs is controlled by the RNA-dependent RNA polymerase Rdr2 but not Rdr1. It will be interesting in the future to determine whether some of these siRNAs carry 5' triphosphate ends. While Rdr1 plays a role in the dsRNA-induced RNAi pathway, Rdr2 was previously shown to be required for all known types of siRNAs in *Paramecium*, including endogenous siRNAs from constitutive SRCs (Carradec et al., 2015; Karunanithi et al., 2019). The biological roles of the 23-nt siRNAs produced from the developmental-specific SRCs and of Rdr2 remain unclear. Further analyses will be needed to determine whether, as hypothesized in Solberg et al., 2023, developmental-specific SRCs regulate aberrant, untranslated mRNAs during development, or instead play a role in other processes such as gene copy number control during development, or act as an antiviral defense mechanism.

**References**

Aury, J.M., Jaillon, O., Duret, L., Noel, B., Jubin, C., Porcel, B.M., Segurens, B., Daubin, V., Anthouard, V., Aiach, N., Arnaiz, O., Billaut, A., Beisson, J., Blanc, I., Bouhouche, K., Camara, F., Duharcourt, S., Guigo, R., Gogendeau, D., Katinka, M., Keller, A.M., Kissmehl, R., Klotz, C., Koll, F., Le Mouel, A., Lepere, G., Malinsky, S., Nowacki, M., Nowak, J.K., Plattner, H., Poulain, J., Ruiz, F., Serrano, V., Zagulski, M., Dessen, P., Betermier, M., Weissenbach, J., Scarpelli, C., Schachter, V., Sperling, L., Meyer, E., Cohen, J., Wincker, P., 2006. Global trends of whole-genome duplications revealed by the ciliate Paramecium tetraurelia. Nature 444, 171–8.

Carradec, Q., Gotz, U., Arnaiz, O., Pouch, J., Simon, M., Meyer, E., Marker, S., 2015. Primary and secondary siRNA synthesis triggered by RNAs from food bacteria in the ciliate Paramecium tetraurelia. Nucleic Acids Research 43, 1818–1833. https://doi.org/10.1093/nar/gku1331

Karunanithi, S., Oruganti, V., Marker, S., Rodriguez-Viana, A.M., Drews, F., Pirritano, M., Nordström, K., Simon, M., Schulz, M.H., 2019. Exogenous RNAi mechanisms contribute to transcriptome adaptation by phased siRNA clusters in Paramecium. Nucleic Acids Res 47, 8036–8049. https://doi.org/10.1093/nar/gkz553

Lepere, G., Nowacki, M., Serrano, V., Gout, J.F., Guglielmi, G., Duharcourt, S., Meyer, E., 2009. Silencing-associated and meiosis-specific small RNA pathways in Paramecium tetraurelia. Nucleic Acids Res 37, 903–15.

Lhuillier-Akakpo, M., Frapporti, A., Denby Wilkes, C., Matelot, M., Vervoort, M., Sperling, L., Duharcourt, S., 2014. Local effect of enhancer of zeste-like reveals cooperation of epigenetic and cis-acting determinants for zygotic genome rearrangements. PLoS Genet. 10, e1004665. https://doi.org/10.1371/journal.pgen.1004665

Marker, S., Carradec, Q., Tanty, V., Arnaiz, O., Meyer, E., 2014. A forward genetic screen reveals essential and non-essential RNAi factors in Paramecium tetraurelia. Nucl. Acids Res. 42, 7268–7280. https://doi.org/10.1093/nar/gku223

Sandoval, P.Y., Swart, E.C., Arambasic, M., Nowacki, M., 2014. Functional Diversification of Dicer-like Proteins and Small RNAs Required for Genome Sculpting. Dev. Cell 28, 174–188. https://doi.org/10.1016/j.devcel.2013.12.010

Sellis, D., Guérin, F., Arnaiz, O., Pett, W., Lerat, E., Boggetto, N., Krenek, S., Berendonk, T., Couloux, A., Aury, J.-M., Labadie, K., Malinsky, S., Bhullar, S., Meyer, E., Sperling, L., Duret, L., Duharcourt, S., 2021. Massive colonization of protein-coding exons by selfish genetic elements in Paramecium germline genomes. PLOS Biology 19, e3001309. https://doi.org/10.1371/journal.pbio.3001309

Solberg, T., Mason, V., Wang, C., Nowacki, M., 2023. Developmental mRNA clearance by PIWI-bound endo-siRNAs in Paramecium. Cell Rep 42, 112213. https://doi.org/10.1016/j.celrep.2023.112213

Zangarelli, C., Arnaiz, O., Bourge, M., Gorrichon, K., Jaszczyszyn, Y., Mathy, N., Escoriza, L., Betermier, M., Regnier, V., 2022. Developmental timing of programmed DNA elimination in Paramecium tetraurelia recapitulates germline transposon evolutionary dynamics. Genome Res. gr.277027.122. https://doi.org/10.1101/gr.277027.122
